# Topology control by a conserved cysteine pair in the OMM-protein CCDC127 enables MICOS interaction

**DOI:** 10.64898/2025.12.12.693940

**Authors:** Christine Zarges, Pauline Schepsky, Robin Alexander Rothemann, Tobias Bock-Bierbaum, Alexander von der Malsburg, Linda Baumann, Aleksandra Trifunovic, Oliver Daumke, Martin van der Laan, Karina von der Malsburg, Jan Riemer

**Author notes:** address correspondence to J.R., K.v.d.M.

## Abstract

Mitochondrial disulfide relay substrates beyond the canonical substrates remain incompletely defined. Revisiting the human MIA40 interactome with enhanced depth, we identified CCDC127 as a previously unrecognized substrate candidate. CCDC127 contains a single transmembrane segment and a conserved C-terminal helical bundle domain (CHB). Comprehensive proteomic and biochemical analyses revealed that, contrary to earlier reports, CCDC127 adopts an N_out_–C_in_ topology in the outer mitochondrial membrane (OMM) with its CHB residing in the intermembrane space (IMS). CCDC127 undergoes oxidation by the disulfide relay, forming a long-range intramolecular disulfide bond between C174 and C219. Loss of these cysteines disrupts correct OMM insertion, inverts transmembrane topology and triggers proteasome-dependent degradation, establishing the disulfide as a key determinant of CCDC127 maturation. Interactome analyses identified MICOS components—particularly the MIC60/MIC19 module—as major partner proteins required for the stability of large oligomeric CCDC127 complexes. CCDC127 deficiency impaired cellular proliferation, influenced phospholipid levels, and caused grossly altered cristae morphology. Together, CCDC127 emerges as a MICOS-associated OMM protein essential for mitochondrial membrane organization and lipid homeostasis.

## INTRODUCTION

The outer membrane of mitochondria (OMM) forms the interface between the cytosol and mitochondria. It plays essential roles in mitochondrial protein import, and in protein and organellar quality control, in the exchange of metabolites, in mitochondrial dynamics and movement, in mediating contacts to other organelles and the inner membrane (IMM), and in signalling the functional state of mitochondria (Becker & Wagner, 2018; Chen *et al*, 2025; Endo & Wiedemann, 2025; Giacomello *et al*, 2020; Helle *et al*, 2013; Mertins *et al*, 2014; Pfanner *et al*, 2019; Tiku *et al*, 2020; Xian & Liou, 2021). Proteins embedded in the OMM are pivotal for its diverse functions. These proteins exhibit various topologies and structural features tailored to their specific roles, which include single- and multi-pass transmembrane domains, β-barrel architectures, and peripheral membrane associations (often tethered to the OMM through amphipathic helices or lipid modifications) (Gupta & Becker, 2021; Rapaport, 2003). Single transmembrane proteins are divided according to their topology: signal-anchored proteins contain an N-terminal transmembrane domain (TMD) and tail-anchored proteins a C-terminal one (Fritz *et al*, 2001; Shore *et al*, 1995; Wattenberg & Lithgow, 2001). In both cases, the bulk of the protein usually faces the cytosol. Moreover, proteins with a single-span internal membrane anchor have been described.

OMM proteins are synthesized in the cytosol and subsequently imported into mitochondria via specialized machineries (Endo & Wiedemann, 2025; Ganesan *et al*, 2024; Gupta & Becker, 2021; Shore *et al*., 1995). For β-barrel proteins, such as porins/VDACs or TOM40, the translocase of the outer membrane (TOM) complex acts as the mitochondrial entry gate. After passage through the TOM complex, additional assistance is provided by small TIM chaperones in the IMS together with the sorting and assembly machinery (SAM) complex and to fold and insert β-barrel proteins into the OMM (Diederichs *et al*, 2020; Ganesan *et al*., 2024; Jores *et al*, 2016; Jores *et al*, 2018; Krimmer *et al*, 2001; Paschen *et al*, 2003; Weinhaupl *et al*, 2018; Wiedemann *et al*, 2003). For multi-pass transmembrane proteins, like yeast Ugo1, a first interaction takes place with Tom70 at the mitochondrial surface from where substrates are passed on to the mitochondrial import (MIM) complex that inserts them into the OMM. Negatively-charged phospholipids like phosphatidic acid and cardiolipin further stimulate the MIM-dependent import of multi-pass transmembrane proteins (Otera *et al*, 2007; Papic *et al*, 2011). Single-span transmembrane proteins such as Mcr1 or Msp1 also rely on the MIM complex or, in human cells, likely on the mitochondrial carrier homolog 2 (MTCH2) (Doan *et al*, 2020; Guna *et al*, 2022; Vitali *et al*, 2020). Single-span transmembrane proteins with larger IMS domains follow more complex routes involving interactions with machineries such as the TIM23 complex (Lauffer *et al*, 2012; Song *et al*, 2014; Wenz *et al*, 2014) (Resch *et al*, 2025) (Sinzel *et al*, 2016)

MM proteins also form direct physical contacts to IMM protein complexes, like MICOS (mitochondrial contact site and cristae organizing system) (Casler & Lackner, 2025; Daumke & van der Laan, 2025; Jenkins *et al*, 2024; Mukherjee *et al*, 2021). While MICOS is particularly important for maintaining the mitochondrial ultrastructure through its role in the formation and stabilization of cristae junctions at the IMM, it also plays pivotal roles in OM-related processes. These include well characterized functions in mitochondrial protein import, *e.g.* through interaction with the TOM, SAM, TIM22 and mitochondrial disulfide relay machineries (Bohnert *et al*, 2012; Callegari *et al*, 2019; Kaurov *et al*, 2018; Varabyova *et al*, 2013; von der Malsburg *et al*, 2011), and in the trafficking of phospholipids (Aaltonen *et al*, 2016; Michaud *et al*, 2016; Monteiro-Cardoso *et al*, 2022). Further studies indicate that MICOS-mediated intra-mitochondrial membrane contact sites may be involved in the regulation of metabolite transport, mitochondrial fusion and fission, and the integration of cellular signalling pathways. Therefore, MICOS is considered as a central hub in mitochondrial and cellular homeostasis (Casler & Lackner, 2025; Daumke & van der Laan, 2025; Jenkins *et al*., 2024; Mukherjee *et al*., 2021).

Here, we characterized the mitochondrial import, maturation, and function of the human OMM protein CCDC127 that we identified as an interaction partner of the mitochondrial disulfide relay system with its core component MIA40 (also known as CHCHD4). The disulfide relay machinery was initially identified in yeast as a specific sorting route for small, soluble cysteine-containing proteins into the IMS (Allen *et al*, 2005; Chacinska *et al*, 2004; Gabriel *et al*, 2007; Mesecke *et al*, 2005; Naoe *et al*, 2004; Terziyska *et al*, 2005). Subsequent studies have substantially extended its substrate spectrum (Habich *et al*, 2019; Kloppel *et al*, 2011; Lionaki *et al*, 2010; Longen *et al*, 2009; Petrungaro *et al*, 2015; Rothemann *et al*, 2024; Terziyska *et al*, 2007; Weckbecker *et al*, 2012; Wrobel *et al*, 2013; Zhuang *et al*, 2013), and work in mammalian cells identified further pathway components and differences in regulation linking disulfide relay-mediated protein import to metabolism (Brosey *et al*, 2025; Fischer *et al*, 2013; Hangen *et al*, 2015; Meyer *et al*, 2015; Rothemann *et al*, 2025; Salscheider *et al*, 2022; Schildhauer *et al*, 2025). CCDC127 presents interesting structural features and was recently suggested to control phospholipid transfer between mitochondria and lipid droplets (Xia *et al*, 2023). We demonstrate for CCDC127 a topology with an N-terminal TMD in the OMM and a large soluble domain in the IMS. We show that MIA40-mediated disulfide insertion into CCDC127 facilitates its translocation into the IMS, which in turn is crucial for acquisition of CCDC127’s correct transmembrane topology, which licenses its association with the MICOS complex. Of note, ablation of MICOS components reduced the steady state protein levels of CCDC127 in mitochondria indicative of an important structural and/or functional connection between MICOS in the IMM and CCDC127 in the OMM. Absence of CCDC127 induces cellular growth defects, results in abnormal mitochondrial cristae morphology and lowered levels of key membrane lipids in particular subfractions of cardiolipin (CL), phosphatidylserine (PS) and phosphatidic acid (PA). Based on our findings, we hypothesize that CCDC127 contributes to the structural organisation and functionality of mitochondria by contributing to membrane lipid homeostasis.

## RESULTS

### CCDC127 is an OMM protein interacting with MIA40

MIA40 loss affects different mitochondrial functions through its activity in the import and folding of IMS proteins. Initial proteomics screens for human MIA40 interaction partners date back more than ten years (Petrungaro *et al*., 2015), and we decided to revisit the MIA40 interactome with unpreceded depth. By combining proteomics approaches with streptactin-based affinity purification of MIA40-Strep from HEK293 cell lines stably and inducibly expressing the protein, we identified 23 previously described MIA40 interaction partners and further 18 proteins that had not been associated with MIA40 before (**Figure 1A**). Among them was CCDC127, a protein with two highly conserved cysteine residues (**Figure 1B, S1A**). We confirmed its interaction with MIA40 in an inverse proteomics experiment after immunoprecipitation of CCDC127-HA from HEK293 cells (**Figure 1C,S1B**). A further validation experiment precipitating MIA40-Strep or CCDC127-HA under both native and denaturing conditions followed by immunoblotting against the respective other protein implied a covalent interaction as would be expected for a substrate of the mitochondrial disulfide relay machinery (**Figure 1D,E**).

**Figure 1.**
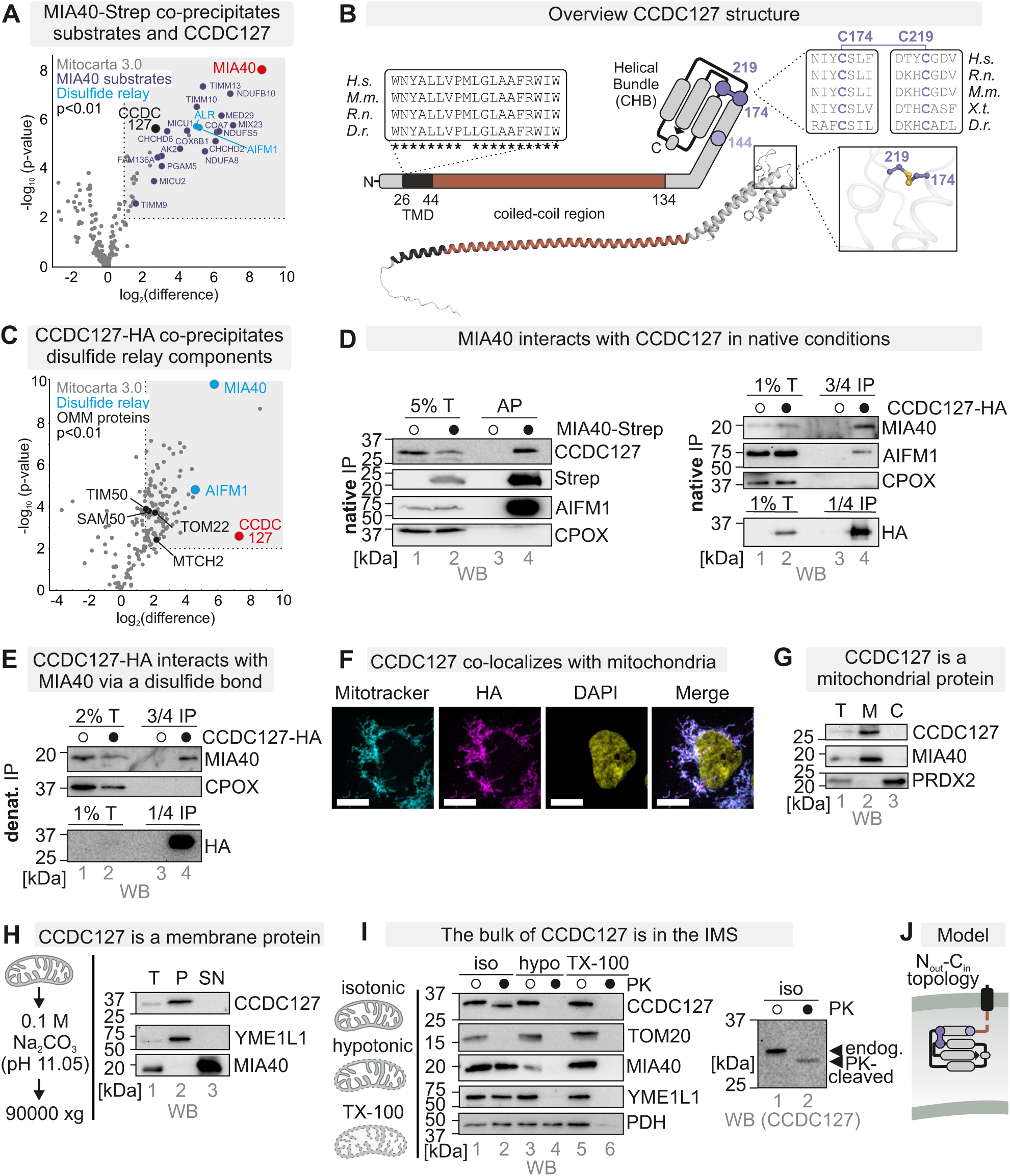
CCDC127 is an OMM protein which interacts with MIA40. **A**: MIA40-Strep co-precipitates CCDC127 in an immunoprecipitation experiment performed after native cell lysis. Cells expressing either MIA40-Strep or the empty vector (Mock) were lysed under native conditions after N-ethyl maleimide (NEM) treatment to prevent further thiol-disulfide exchange. MIA40-Strep was enriched using strep-tactin beads. Eluates were analyzed by quantitative label-free mass spectrometry. AIFM1 and ALR as parts of the mitochondrial disulfide relay are highlighted in blue. Known MIA40 substrates are labeled in lilac. n = 4 biological replicates, an unpaired one-sample two-sided Student’s t-test was applied (P < 0.01, log_2_-enrichment >1). **B.** Schematic representation of the CCDC127 protein. CCDC127 contains a conserved hydrophobic stretch at its N-terminus (TMD, aa 26-44) labelled in black, a coiled-coil region (aa 44-134) colored in brown and a C-terminal helical bundle domain (CHB, aa 134-260) domain with three cysteine residues, of which two are conserved and predicted to form a disulfide bond according to the AlphaFold3 model (C174, C219). **C.** CCDC127-HA co-precipitates AIFM1 and MIA40 under native conditions. Cells expressing either CCDC127-HA, or the empty vector (Mock) were lysed under native conditions and CCDC127-HA was enriched using HA-antibody beads. Eluates were analyzed by quantitative label-free mass spectrometry. AIFM1 as part of the mitochondrial disulfide relay as well as different OMM proteins are highlighted. n = 4 biological replicates, an unpaired one-sample two-sided Student’s t-test was applied (P < 0.01, log_2_-enrichment >1.5). **D.** MIA40 interacts with CCDC127 under native conditions. Cells expressing either MIA40-Strep (*left*) or CCDC127-HA (*right*) were lysed under native conditions after treatment with N-ethylmaleimide (NEM). Proteins were precipitated from the lysates using strep-tactin beads or HA-antibody beads, respectively. AIFM1 is part of the disulfide relay system and is co-precipitated with MIA40-Strep and CCDC127-HA. The IMS protein CPOX served as a non-interacting specificity control. **E.** CCDC127-HA interacts with MIA40 under denaturing conditions via a mixed disulfide bond. Cells expressing CCDC127-HA were lysed under denaturing conditions after treatment with N-ethylmaleimide (NEM). Proteins were precipitated from the lysates using HA-antibody beads, respectively. The IMS protein CPOX served as a non-interacting specificity control. **F.** Immunofluorescence of CCDC127-HA shows co-localization with Mitotracker. Cells expressing CCDC127-HA were fixed, permeabilized, and stained using a primary antibody against the HA epitope or Mitotracker. Cells were analysed by fluorescence microscopy. Mitotracker signal served as positive control for mitochondria and DAPI as marker for the nucleus. Magenta, HA-signal; Yellow, DAPI signal; Cyan, mitotracker signal. A white colour indicates signal overlap. Bar corresponds to 10 μm. n = 2 biological replicates. **G.** CCDC127 is a mitochondrial protein. Cells were lysed by potter homogenization followed by centrifugation to separate a mitochondrial (“M”) and a post-mitochondrial (“C”) fraction. The total lysate (“T”) served as loading control, and MIA40 and PRDX2 as controls for mitochondria and cytosol, respectively. n = 3 replicates. **H.** CCDC127 is a membrane protein. Mitochondria isolated from HEK293 cells were treated with Na_2_CO_3_, pH 11.05. Then supernatant and pellet were separated by ultracentrifugation. MIA40 and YME1L served as soluble and membrane controls, respectively. n = 2 replicates. **I.** CCDC127 is an OMM protein with the major portion of the protein localized in the IMS. Mitochondria were enriched and then either incubated in an isotonic buffer (iso), in a hypotonic buffer (hypo) or a buffer containing Triton X-100 (TX-100). The resulting fractions were either treated with proteinase K (PK) to test for protection of proteins or left untreated. TOM20, MIA40, YME1L and PDH served as OMM, IMS, IMM and matrix controls, respectively. PK treatment of intact mitochondria (*lane 2*) resulted in partial processing of the entire pool of CCDC127 indicating a small portion of the protein to be exposed to the cytosol. n = 2 replicates. **J.** Model indicating the N_out_-C_in_ topology of CCDC127 in the OMM.

CCDC127 is a protein of 260 amino acids that carries a conserved hydrophobic stretch close to its N-terminus likely serving as a transmembrane anchor (**Figure 1B**). This potential TMD is followed by an almost 100-amino acid-long region that is proposed to form a coiled-coil domain in an AlphaFold 3 structural model. At the C-terminus of CCDC127 a helical bundle domain (CHB) is predicted. It contains two conserved cysteines (C174, C219) that are proposed to form a disulfide bond and a third less conserved cysteine (C144). Notably, CCDC127 does not appear to carry a classical mitochondrial targeting signal.

CCDC127 has recently been described to localize to the OMM with the bulk of the protein including its conserved cysteine residues facing the cytosol (Xia *et al*., 2023). Because such a topology would exclude a disulfide-mediated interaction with MIA40, we revisited the localization and topology of CCDC127. By combining immunofluorescence analysis, cellular fractionation approaches, and carbonate extraction assays, we confirmed mitochondrial and membrane localization of CCDC127 (**Figures 1F-H**). However, Proteinase K (PK) treatment of isolated mitochondria revealed that CCDC127 was largely protected in intact mitochondria (**Figure 1I***, lane 2*). Upon opening of the OMM using hypo-osmotic conditions, this PK protection was lost indicating an IMS localization of the bulk of the protein (**Figure 1I***, lane 4*). A closer look at CCDC127 in intact mitochondria treated with PK or left untreated confirmed the initial impression that CCDC127 slightly truncated under PK-treated conditions suggesting that a small part of CCDC127 is exposed to the cytosol (**Figure 1I**, *right zoom-in*). To precisely determine the topology of CCDC127 in the OMM, we employed CCDC127 variants that were either C-terminally HA or N-terminally FLAG tagged (**Figure S1B,C**). Both variants were localized to mitochondria and were inserted into the OMM (**Figure S1D-G**). PK treatment of intact mitochondria bearing either of the differentially tagged CCDC127 variants revealed that the C-terminally fused HA-epitope was protected from PK digestion, whereas the N-terminal FLAG-tag was not (**Figure S1H,I**). Based on this comprehensive biochemical analysis, we propose a new N_out_-C_in_-topology for the CCDC127 protein with a TMD close to the N-terminus inserting it into the OMM and a large hydrophilic portion including the CHB domain that is localized to the IMS (**Figure 1J**). The positioning of the TMD in the OMM is also in line with the recovery of various OMM proteins in the CCDC127 coprecipitation analysis (**Figure 1C**).

### The CCDC127 TMD enables targeting to the OMM

We next assessed the role of the different parts of CCDC127 for its unusual localization and topology. To this end, we generated a series of stable cell lines expressing truncation variants (**Figure 2A**): one lacking the cytosolic part of the protein (CCDC127^24-260^), one lacking the cytosolic part and the TMD (CCDC127^45-260^), one only containing the CHB (CCDC127^135-260^), and one lacking this region (CCDC127^1-134^).

**Figure 2.**
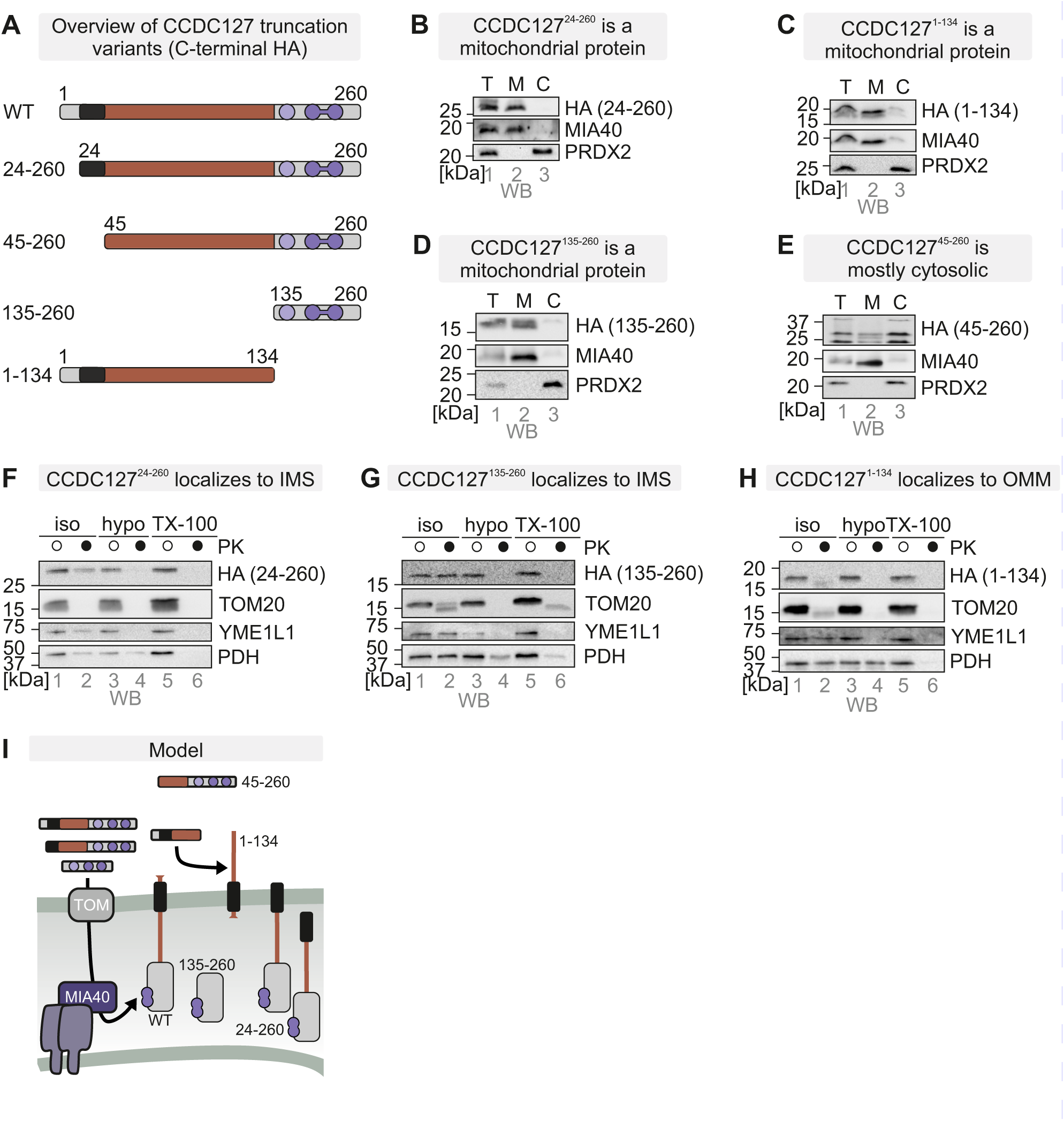
The CCDC127 TMD enables targeting to the OMM. **A.** Overview of CCDC127 truncation variants with C-terminal HA tag to analyze determinants for OMM topology and localization. **B-E.** CCDC127^24-260^ **(B)**, CCDC127^1-134^ **(C)**, CCDC127^135-260^ **(D)**, truncation variants localize to mitochondria whereas CCDC127^45-260^ **(E)** localizes mostly to the cytosol. Cells were lysed by potter homogenization followed by centrifugation to separate a mitochondrial (“M”) and a post-mitochondrial (“C”) fraction. The total lysate (“T”) served as loading control, and MIA40 and PRDX2 as marker for mitochondria and cytosol, respectively. **F.** Truncation of the cytosolic amino acids 1-23 does not affect IMS localization of CCDC127. Experiment was performed as in **Figure 1I** except with cell lines expressing CCDC127^24-260^-HA. n = 3 replicates. Iso, mitochondria treated with isotonic buffer; hypo, mitochondria treated with hypo-osmotic buffer. **G.** The CHB of CCDC127 is sufficient for IMS localization of CCDC127. Experiment was performed as in **Figure 1I** except with cell lines expressing CCDC127^135-260^-HA. n = 2 replicates. Iso, mitochondria treated with isotonic buffer; hypo, mitochondria treated with hypo-osmotic buffer. **H.** A CCDC127 variant lacking the CHB is partially exposed to the cytosol. Experiment was performed as in **Figure 1I** except with cell lines expressing CCDC127^1-134^-HA. n = 2 replicates. Iso, mitochondria treated with isotonic buffer; hypo, mitochondria treated with hypo-osmotic buffer. **I.** Model depicting the necessity of TMD for mitochondrial CCDC127 import and membrane insertion.

CCDC127^24-260^, CCDC127^1-134^ and CCDC127^135-260^ still localized to mitochondria, while CCDC127^45-260^ localized to the cytosol (**Figure 2B-E, S2A**) implying that the transmembrane domain was important for mitochondrial localization of CCDC127 variants. An exception was the CHB domain that if expressed without other parts of CCDC127 was able to drive mitochondrial localization. Notably, the CCDC127^45-260^ variant appeared to undergo some processing as it always was present in a double band.

By mitochondrial subfractionation, we found CCDC127^24-260^ and CCDC127^135-260^ to be protected from PK (**Figure 2F,G**), whereas CCDC127^1-134^ was PK-accessible (**Figure 2H**). Together with the finding that CCDC127^1-134^ still behaved as a membrane protein (**Figure S2B**) this implied a localization of this variant to the cytosolic face of the OMM. As expected, the variant lacking TMD and the coiled-coil region (CCDC127^135-260^) behaved like a soluble IMS protein, while the TMD-containing CCDC127^24-260^ was partially soluble suggesting some role for the small cytosolic stretch in CCDC127 for correct and efficient OMM insertion (**Figure S2C-E**). Collectively, we found that the TMD drives OMM insertion of CCDC127 and the CHB determines the N_out_-C_in_ topology by guiding the protein into the IMS. Presence of the coiled-coil region in the absence of the TMD seems to hamper translocation into the IMS (**Figure 2I**).

### The disulfide relay oxidizes CCDC127 during its maturation

The interaction with MIA40 suggests that CCDC127 acquires a disulfide bond through the mitochondrial disulfide relay. To experimentally test for the presence of this disulfide, we employed a maleimide shift assay in which reduced but not oxidized cysteine residues are modified with the maleimide mmPEG_24_ thereby increasing the molecular weight of the modified protein (**Figure 3A**). As control for the maximal possible migration shift (representing reduced protein), we used cells that were lysed and completely reduced under denaturing conditions and then incubated with mmPEG_24_. As minimal unshifted control we employed untreated (representing oxidized protein) cell lysate analyzed by reducing SDS-PAGE. CCDC127 migrated close to the height of the untreated control indicating it to be oxidized at steady state (**Figure 3B**, *lane 2*). In this experiment, we also found no indications for a disulfide-linked dimer of CCDC127 which would migrate at around 65-75 kDa in lane 2.

**Figure 3.**
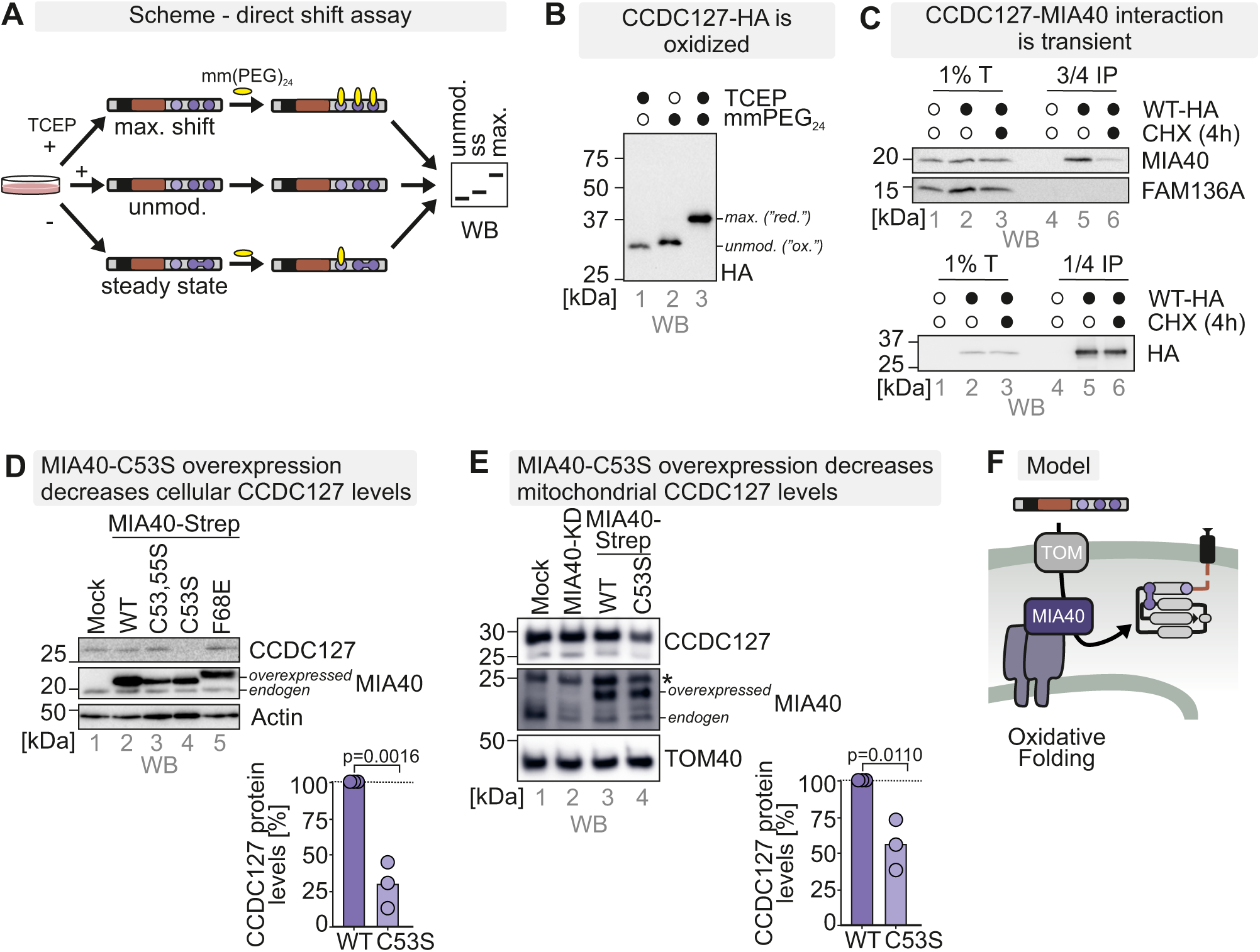
The disulfide relay oxidizes CCDC127 during its maturation. **A.** Direct redox shift assay to assess protein thiol redox states. Cells were lysed and treated with the maleimide mmPEG_24_ that modifies free thiols (reduced) but not thiols in disulfide bonds (oxidized). Modification of proteins with mmPEG_24_ results in a slower migration of the protein on SDS-PAGE. Unmodified cell lysates (unmod.) and cell lysates pretreated with the reductant TCEP (max. shift) served as controls. Ss indicates the steady state redox state of the protein. **B.** Cysteines in CCDC127 are present in the oxidized state. As described in (**A**). Cells were lysed and either treated with the strong reductant TCEP (*lanes 1 and 3*) or left untreated (*lane 2*). Then lysates were left untreated (*lane 1, unmod.*) or incubated with mmPEG_24_ (*lanes 2, ss and 3, max. shift*). Lysates were analyzed by SDS-PAGE and immunoblotting. Cysteines in CCDC127-HA are present in the oxidized state. n = 3 replicates. **C.** CCDC127-MIA40 interaction is transient. Cells expressing CCDC127-HA were untreated or treated with 100 µg/ml cycloheximide (CHX) for 4 h prior to cell lysis under native conditions after treatment with N-ethylmaleimide. Interaction partners of CCDC127-HA were enriched using HA-antibody beads. The IMS protein FAM136A served as a non-interacting specificity control. **D.,E.** Overexpression of the dominant-negative MIA40^C53S^ mutant decreases cellular (**D**) and mitochondrial (**E**) CCDC127 levels. Expression of MIA40 variants lacking either both cysteines of the redox active CPC motif (C53S,C55S) or only C53 (C53S; “dominant-negative variant”) or with a mutation in the chaperone site of MIA40 (F68E) was induced for 5 days in glucose-containing medium. Mitochondria were isolated and lysed, or cells were directly lysed, and protein levels were analyzed by SDS-PAGE and immunoblot against the indicated proteins. n = 3 replicates (one sample t-test). **F.** Model depicting the necessity of disulfide bond formation for mitochondrial CCDC127 import.

Next, we tested whether the interaction between CCDC127 and MIA40 was transient as this would suggest that MIA40 introduces disulfide bonds into CCDC127 during import and maturation. To this end, we incubated cells with the translation inhibitor cycloheximide and probed then for the MIA40-CCDC127 interaction. A transient but not a stable interaction would be affected by the cycloheximide treatment. Indeed, upon treatment of cells with cycloheximide, we lost the MIA40-CCDC127 interaction suggesting a transient interaction between MIA40 and CCDC127 during CCDC127 import to introduce a disulfide bond into the protein (**Figure 3C**).

Often protein levels of disulfide relay substrates are affected by MIA40 or AIFM1 loss or the overexpression of a redox-inactive dominant-negative C53S variant of MIA40. Interestingly, CCDC127 levels in whole cells were not affected by the loss of MIA40 or AIFM1 (**Figure S3**), but by the expression of MIA40^C53S^ (**Figure 3D**, *lane 4*). MIA40^C53S^ expression also led to reduced CCDC127 amounts in isolated mitochondria (**Figure 3E**, *lane 4*).

Collectively, we demonstrated that MIA40 forms a disulfide bond in CCDC127. This disulfide bond is unusual for MIA40 substrates because it is a long-range disulfide bond connecting C174 and C219, and because it does not connect two cysteines situated in α-helices as C219 is positioned in a loop and C174 at the very end of an α-helix (**Figure 3F**). The localization of the CHB alone to the IMS (**Figure 2G**) further suggests that the disulfide relay is capable of determining IMS localization of this CCDC127 domain.

### Cysteines in CCDC127 control its orientation in the OMM

To address the role of the CCDC127 cysteines for the topology of the overall protein, we generated cell lines stably expressing cysteine-to-alanine variants: CCDC127^C174A, C219A^ (2CA), CCDC127 ^C144A, C174A, C219A^ (3CA), and CCDC127^C144A^ (**Figure 4A**). Notably, the 2CA and the 3CA variants of CCDC127 exhibited strongly lowered cellular protein levels indicating the importance of the disulfide bond for the stability of CCDC127 (**Figure 4B**)

**Figure 4.**
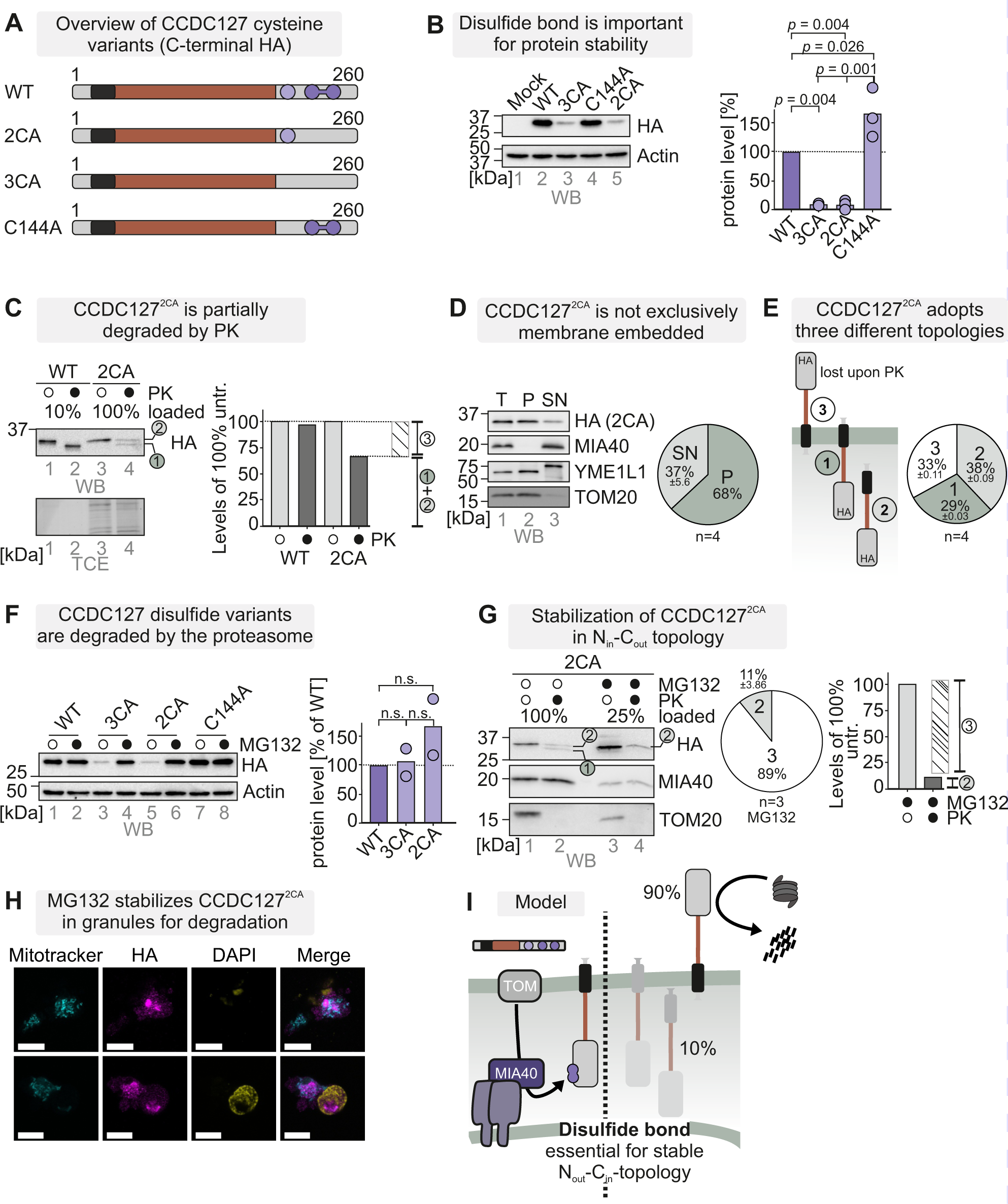
Cysteines in CCDC127 control its orientation in the OMM. **A.** Overview of CCDC127 cysteine variants for analysis of determinants of OMM topology and localization. **B.** CCDC127 variants lacking the disulfide-forming cysteines are unstable. Lysates from cells expressing cysteine variants of CCDC127-HA (2CA: C174A, C219A, 3CA:C144A, C174A, C219A) were analyzed by reducing SDS-PAGE and immunoblotting. Signals were quantified using ImageLab, and the amount of protein was plotted. N = 3 replicates (one-way ANOVA with post hoc Tukey HSD test). **C.** The CCDC127-2CA-HA variant is only partially protected from proteinase K (PK) treatment in intact mitochondria. Experiment was performed as in Figure 1I except with cell lines expressing CCDC127-2CA-HA. Signals were quantified using ImageLab, and the percentage of fractions was plotted. **D.** The CCDC127-2CA-HA variant is only partially inserted into the membrane. Experiment was performed as in Figure 1H except with cell lines expressing CCDC127-2CA-HA. Signals were quantified using ImageLab, and the percentage of soluble and pellet (membrane) fraction was plotted. n = 4 replicates. T, total; P, pellet (membrane), SN, soluble. **E.** CCDC127-2CA-HA variant adopts three different topologies. Fraction #1 represents a WT-like behavior, fraction #2 represents a PK-protected IMS-localized, soluble CCDC127. The difference between PK-treated and -untreated signal represents a largely cytosol-exposed fraction #3. n = 4 replicates. **F.** CCDC127 variants lacking the disulfide-forming cysteines are degraded by the proteasome. Lysates from cells expressing cysteine variants of CCDC127-HA and treated with MG132 or left untreated were analyzed by reducing SDS-PAGE and immunoblotting. Signals were quantified using ImageLab, and the amount of protein was plotted. n = 3 (-MG132) N = 2 (+MG132) replicates (one-way ANOVA with post hoc Tukey HSD test). **G.** The CCDC127-2CA-HA variant is only partially protected from proteinase K (PK) treatment in intact mitochondria after 16h MG132 treatment. Inhibition of the proteasome stabilizes CCDC127-2CA in the N_in_-C_out_-topology, which is inverse from the WT (*i.e.* the PK-sensitive fraction #3 is very large). Signals were quantified using ImageLab, and the percentage of fractions was plotted. n = 3 replicates. **H.** The CCDC127-2CA-HA variant is stabilized by MG132 in granules for degradation. Immunofluorescence of CCDC127-2CA-HA after 16h MG132 treatment was performed as in **Figure 1F**. Bar corresponds to 10 μm. n = 2 replicates. **I.** Model indicating that the CHB of CCDC127 and its disulfide-forming cysteines are critical for the correct orientation in the OMM and the stability of CCDC127.

We then subjected isolated mitochondria containing the 2CA variant to a submitochondrial fractionation assay (**Figure 4C**, *2CA - loaded ten times more compared to WT*). By comparison to WT CCDC127, we found that the residual amounts of the 2CA variant were reproducibly protected to about 2/3 from PK treatment in intact mitochondria (**Figure 4C, S4A**, *fractions 1+2*) indicating a partial localization of the protein’s soluble domain to the cytosolic face of the OMM. Moreover, PK digest yielded a protected double band for the 2CA variant indicating that about half of the PK-protected protein (*i.e.* 1/3 of the total protein) was not membrane-inserted into the OMM (**Figure 4C***, fraction 2*). In line, we found the 2CA variant to be partially soluble in a carbonate extraction experiment (**Figure 4D**). Based on this data, we propose that only a part of the 2CA variant reaches the IMS, and that its insertion from the IMS into the OMM is hampered (**Figure 4E**).

Proteasome inhibition by MG132 strongly stabilized the 2CA variant (**Figure 4F**). Moreover, a large fraction of CCDC127-2CA became sensitive to PK treatment in mitochondria isolated from these MG132-treated cells (**Figure 4G**, *lane 4*). This indicates that the 2CA variant is largely inserted into the OMM in a non-native N_in_-C_out_-topology (∼90%). As a consequence, it is likely continuously extracted and then degraded by the proteasome system.

Interestingly, while the C144A variant colocalized with a mitochondrial marker in immunofluorescence, both the 2CA and the 3CA variant exhibited a punctuated stain pattern that only partially overlapped with mitochondria in immunofluorescence imaging (**Figure S4B**). This became even more pronounced upon proteasomal inhibition (**Figure 4H**) suggesting that cysteine variants of CCDC127 localized to non-mitochondrial structures likely *en route* to degradation.

Collectively, our data are in line with a model, in which the TMD and the CHB with its cysteines are critical determinants of CCDC127 topology (**Figure 4I**). While the TMD controls OMM insertion, variants without cysteines still enter the OMM but face the cytosol with the bulk of the protein instead of the IMS rendering them prone to degradation.

### CCDC127 is a new interactor of the MICOS complex

The intricate topology of CCDC127 together with its coiled-coil and CHB domains made us search for functional interaction partners of the mature protein in the IMS. We went back to our initial CCDC127 interactome screen (**Figure 1C**) and looked through the spectrum of candidate proteins. Of note, several IMM proteins were identified as potential interaction partners suggesting that the extended IMS domain of CCDC127 is in close proximity to the IMM or even directly participates in the formation of intramitochondrial membrane contact sites. In support of this view, the IMS-exposed MICOS components MIC60 and MIC19 were amongst the most prominent hits of our screen (**Figure 5A**). The MIC60/MIC19 module of MICOS is known to interact with the SAM complex of the OMM via the central SAM50 subunit (Ding *et al*, 2015; Harner *et al*, 2011; Tang *et al*, 2020), which we also identified as putative CCDC127 interactor (**Figure 1C**). Also, in line with our findings is the detection of crosslinks between CCDC127 and the MICOS subunits MIC60, MIC19 and MIC25 in a recent high-throughput spatial proteomics study (Zhu *et al*, 2024) (**Figure 5B**). To confirm the CCDC127-MICOS interaction, we generated a cell line expressing C-terminally FLAG-tagged MIC10. We then isolated the MICOS complex and its interacting proteins via immunoprecipitation. Besides known MICOS components, we indeed co-purified substantial amounts of CCDC127 with MIC10-FLAG (**Figure 5C**). Taken together, we have identified CCDC127 as a novel interaction partner of the MICOS complex.

**Figure 5.**
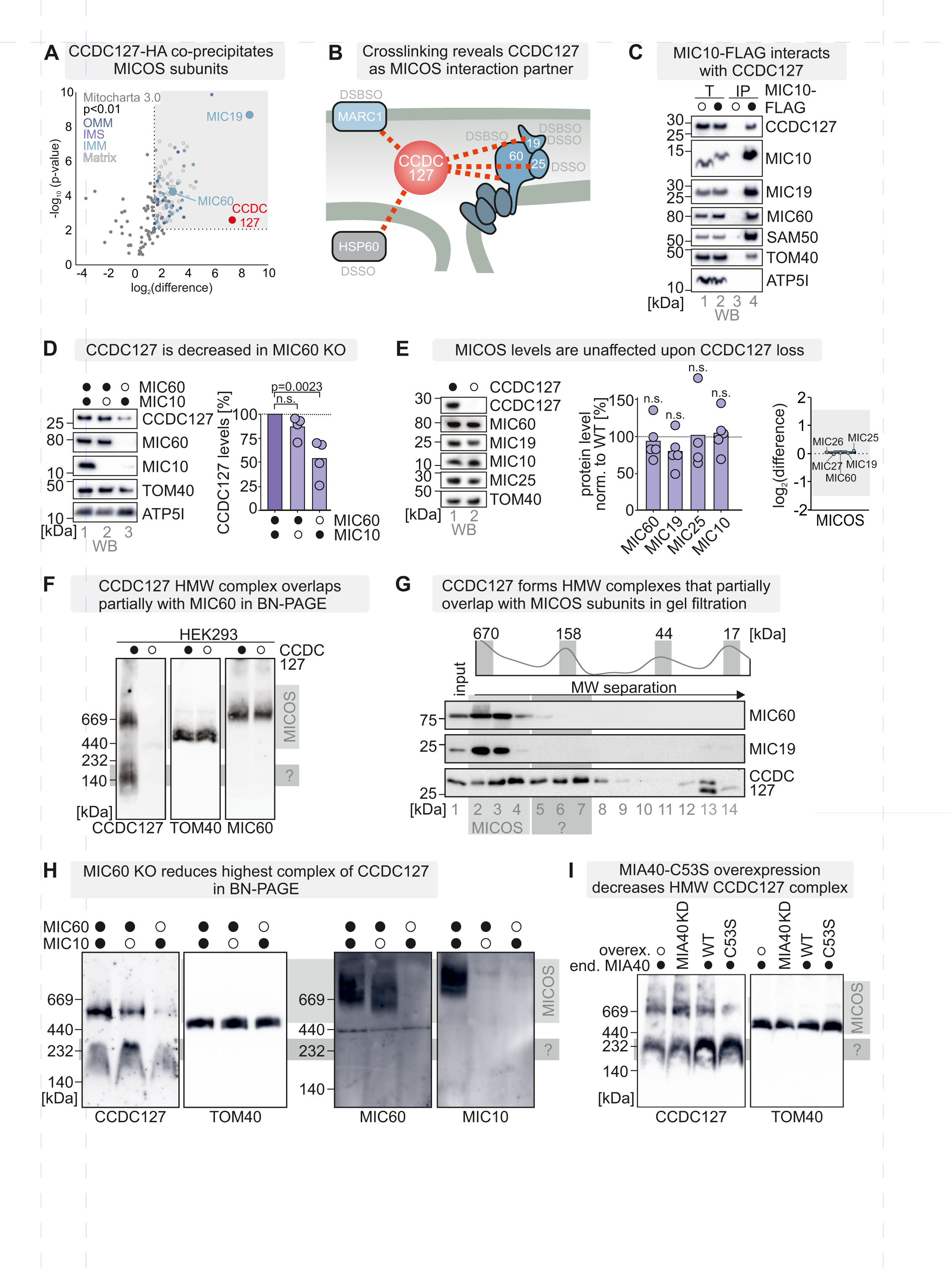
CCDC127 is an interaction partner of the MICOS complex. **A.** CCDC127-HA co-precipitates MIC19 and MIC60 under native conditions. Data are the same as shown in Figure 1C. Cells expressing either CCDC127-HA, or the empty vector (Mock) were lysed under native conditions and CCDC127-HA was enriched using HA-antibody beads. Eluates were analyzed by quantitative label-free mass spectrometry. n = 4 biological replicates, an unpaired one-sample two-sided Student’s t-test was applied (P < 0.01, log_2_-enrichment >1). **B.** Crosslinking assisted spatial proteomics study revealed CCDC127 as MICOS interaction partner. Beside MIC60, MIC19 and MIC25, HSP60 and MARC1 were found as crosslinked partners of CCDC127 in a recently published study (Zhu *et al*., 2024). **C.** MIC10-FLAG interacts with CCDC127 under native conditions. WT mitochondria or mitochondria containing MIC10-FLAG were solubilized in digitonin and MIC10-FLAG was precipitated using FLAG-antibodies. Immunoblot analyses were performed against CCDC127 and MIC10 as well as subunits of the MICOS complex as positive controls and ATP5I as negative control. **D.** CCDC127 protein levels are decreased in MIC60 KO HEK293 mitochondria. Protein levels in isolated mitochondria from HEK293 cell lines depleted of MIC10 (*lane 2*) or MIC60 (*lane 3*) were compared to protein levels in corresponding wild-type (WT) mitochondria. Mitochondria were analyzed by reducing SDS-PAGE and immunoblotting against the indicated proteins. Signals were quantified using ImageLab, and the amount of protein was plotted. n = 4 replicates. TOM40 and ATP5I served as loading control. **E.** MICOS levels are unaffected by CCDC127 depletion. Whole cell proteomes of CCDC127-KO cells. Cells were lysed under native conditions and analyzed by label-free mass spectrometry. N = 4 biological replicates, an unpaired one-sample two-sided Student’s t-test was applied. Protein levels in cells depleted of CCDC127 were compared to protein levels in corresponding wild-type (WT) cells. Lysates were analyzed by reducing SDS-PAGE and immunoblotting against the indicated proteins. Signals were quantified using ImageLab, and the amount of protein was plotted. n = 3 replicates. Pyruvate dehydrogenase (PDH) served as loading control (one-way ANOVA with post hoc Tukey HSD test). **F.** Endogenous CCDC127 is present in two high molecular weight (HMW) complexes. Mitochondria isolated from WT- and CCDC127 KO cells were resolved with blue native electrophoresis (BN–PAGE) followed by immunoblotting against the indicated proteins. The identity of the smaller ∼140 kDa complex remains unclear. **G.** A higher oligomer complex containing endogenous CCDC127 migrates in same similar MW range as the MICOS components MIC60 and MIC19 in gel filtration. HEK293 cells were lysed under native conditions (Triton X-100), and the cleared lysates subjected to gel filtration analysis. Eluted fractions were subjected to TCA precipitation, resuspension in loading buffer containing SDS and DTT, and subsequent immunoblotting against CCDC127, MIC19 and MIC60. Endogenous CCDC127 migrates in two complexes at around 150 kDa and 670 kDa (as judged by comparison to protein markers: apoferritin 443 kDa; β-amylase 200 kDa; alcohol dehydrogenase 150 kDa; bovine serum albumin 66 kDa; carbonic anhydrase 29 kDa). **H.** Levels of the higher 670 kDa CCDC127 complex levels are affected in MIC10 KO and MIC60 KO HEK293 cells. Complex levels in HEK293 cell lines depleted of MIC10 or MIC60 were compared to protein levels in corresponding wild-type (WT) cells. Lysates were analyzed by BN-PAGE. **I.** Levels of the higher 670 kDa CCDC127 complex levels are affected upon MIA40^C53S^ overexpression. Complex levels in HEK293 cell lines expressing the C53S variant of MIA40 were compared to protein levels in corresponding wild-type (WT) cells. Lysates were analyzed by BN-PAGE.

In order to assess the stability and functional importance of the CCDC127-MICOS interaction, we analysed mitochondria isolated from MIC10 and MIC60 KO cell lines for their CCDC127 content. Loss of MIC60 leads to an almost complete loss of the entire MICOS complex (Stephan *et al*, 2020) and also resulted in reduced levels of CCDC127 indicative of an intimate structural or functional connection (**Figure 5D***, lane 3*). Mitochondria lacking MIC10 still accumulate close-to-normal levels of MIC60 and also retained WT-like amounts of CCDC127 (**Figure 5D***, lane 2*). We conclude that the stability of CCDC127 in mitochondria depends on the MIC60/MIC19 module of MICOS and, thus, rather on OMM-IMM contact site formation than on the stability of cristae junctions. Conversely, CCDC127-deficient mitochondria still contained normal levels of MICOS subunits indicating CCDC127 as peripheral partner of the complex (**Figure 5E**).

Both, blue native-PAGE analysis (BN-PAGE, **Figure 5F**) and size-exclusion chromatography (**Figure 5G**) revealed the presence of high-molecular-weight CCDC127 complexes in mitochondria. The largest CCDC127-containing complex of ∼600 kDa shows a similar migration behaviour on BN-PAGE and chromatography profiles as the most abundant MICOS complex form. However, this complex species does not contain CCDC127 and MICOS together, because CCDC127-deficient mitochondria still exhibit normal levels of the respective MICOS complex (**Figure 5F**). We conclude that CCDC127 and MICOS may be part of a large membrane-spanning protein assembly in mitochondria that falls apart into protomers during harsh experimental procedures, like chromatography or electrophoresis. Finally, BN-PAGE analysis of MIC10- and MIC60-deficient mitochondria demonstrated that, primarily, loss of the ∼600 kDa CCDC127 complex accounts for the reduced protein levels, while lower molecular weight species are largely stable (**Figure 5H**). This is also in line with the finding that expression of MIA40^C53S^ that results in lowered CCDC127 levels (**Figure 3E**) leads to the loss of primarily the ∼600 kDa complex (**Figure 5I**).

### Loss of CCDC127 impairs cellular fitness, lipid composition and ultrastructure

MICOS plays important roles in organizing mitochondrial membrane architecture, it is critical for efficient ATP generation, impacts on lipid metabolism and protein import into mitochondria. We wondered whether CCDC127 as MICOS partner protein of the OMM would impact cellular fitness in a similar manner. In our growth assays, we found that proliferation of CCDC127-deficient cells on glucose-containing medium was impaired. This became more pronounced when we shifted cells to galactose-containing medium that requires a fully oxidative phosphorylation machinery (**Figure 6A**).

**Figure 6.**
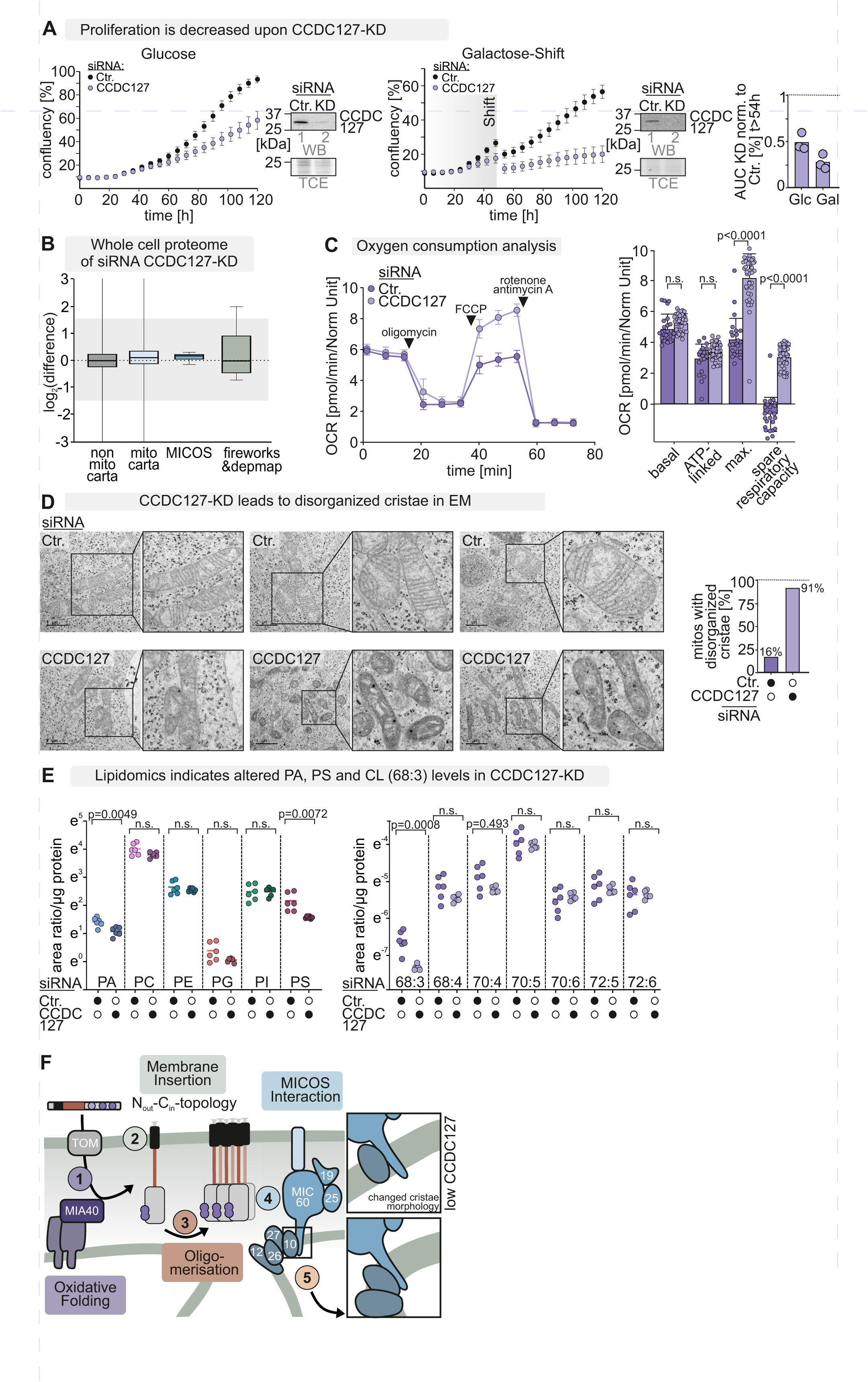
Loss of CCDC127 impairs cellular proliferation, lowers cardiolipin levels and results in abnormal cristae. **A.** Cell proliferation upon acute loss of CCDC127 in HEK293 cells is impaired. Cells were treated with siRNA to deplete CCDC127. Subsequently, their proliferation on glucose-containing medium (*left*) or galactose-containing medium (*right)* was assessed in a high-throughput microscopy assay by automatically scoring their confluency. The depletion of CCDC127 was confirmed after 120 h by SDS-PAGE and immunoblotting. N=3 replicates. **B.** Box plot analysis of whole cell proteomes of cells depleted from CCDC127. Cells were lysed under native conditions and analyzed by label-free mass spectrometry. Indicated protein groups are depicted as box plots. N = 4 biological replicates, an unpaired one-sample two-sided Student’s t-test was applied. Mitocarta 3.0, MICOS and fireworks and depmap hits are represented according to the log2 enrichment. Grey area represents abundance of proteins with minor changes. Further analysis in Figure S5A-C. **C.** Cells lacking CCDC127 show increased maximum respiration. Oxygen consumption profile was analyzed after 72h of siRNA-mediated depletion of CCDC127 using a Seahorse analyser. Basal, ATP-linked and non-mitochondrial respiration were unchanged and maximal respiratory capacity was increased. N=24 (ctr.) n=40 (KD) in two independent experiments. OCR was normalized to cell density. **D.** Depletion of CCDC127 leads to disorganized cristae in transmission electron microscopy (EM). Cells were treated for 72 hours with a control siRNA or siRNA against CCDC127. Cells were fixed and analyzed by EM. Mitochondria with abnormal cristae structures were counted. Three representative images are shown per condition. N=3 replicates. Bar=1µm **E.** Lipidomics analysis of cells depleted of CCDC127. Cells were treated for 72 hours with a control siRNA or siRNA against CCDC127. Cells were pelleted and lipidomics analysis was performed. Levels of phosphatidic acid (PA) and phosphatidylserine (PS) as well as subfractions of cardiolipin (CL) were lowered in cells with CCDC127 loss. Levels of phosphatidylglycerol (PG), phosphatidylcholines (PC), phosphatidylinositol (PI) and phosphatidylethanolamine (PE) were unchanged. Further analysis in **Figure S6**. **F.** Model. CCDC127 is an OMM protein linked to MICOS and controlling lipid homeostasis and mitochondrial ultrastructure. *Step 1*. Upon import into the IMS, CCDC127 acquires a long-range disulfide bond catalyzed by the mitochondrial disulfide relay. *Step 2*. Insertion of CCDC127 into the OMM in an N_out_-C_in_ toplogy is facilitated by its TMD. *Step 3*. CCDC127 homo-oligomers integrate into a 600 kDa complex. *Step 4*. CCDC127 stability depends on its interaction with MICOS components. *Step 5*. CCDC127 depletion affects mitochondrial ultrastructure, and cellular fitness.

Next, we investigated changes in the whole cell proteome upon CCDC127 loss. We found only little changes in the cellular and mitochondrial proteomes, MICOS or in proteins sharing DEPMAP and FIREWORKS profiles with CCDC127 (**Figure 6B, S5A,B**).

We then analyzed the oxygen consumption profile of control- and CCDC127-siRNA treated cells using a Seahorse analyzer that provides insights into the functioning of the respiratory chain. We thereby found unchanged basal, ATP-linked and non-mitochondrial respiration but an increased maximal respiratory capacity (**Figure 6C**). Collectively, this suggests that CCDC127 is not essential for the core electron transport chain but might be involved in a regulatory pathway that can be bypassed or altered.

Finally, we wondered whether CCDC127 is important for the maintenance mitochondrial morphology and membrane architecture. We observed by transmission electron microscopy (TEM) that CCDC127-deficient mitochondria exhibit severely altered cristae morphologies (**Figure 6D**). Other cellular organelles appeared not to be affected by CCDC127 reduced levels in our TEM analysis. Observing these strong differences, we wondered whether the cellular membrane composition was affected and thus performed lipidomics experiments. Levels of phosphatidic acid (PA) and phosphatidylserine (PS) were slightly but significantly lowered in cells upon CCDC127 ablation. Levels of phosphatidylglycerol (PG), phosphatidylcholines (PC), phosphatidylinositol (PI) and phosphatidylethanolamine (PE) were unchanged. However, the amounts of the mitochondrial signature phospholipid cardiolipin were substantially reduced upon loss of CCDC127 (**Figure 6E, S6**).

Collectively, our physiological assays imply a function of CCDC127 in maintaining mitochondrial morphology and membrane lipid homeostasis, in particular cardiolipin levels.

## DISCUSSION

### Disulfide bond formation as regulator of CCDC127 topology

In this study, we identified substrates of the mitochondrial disulfide relay. Among them, we found CCDC127, a protein with an unusual structure and topology for a MIA40 substrate. While the TMD drives the protein to the OMM, its CHB domain bearing the disulfide-forming cysteines is responsible for orientation in a N_out_-C_in_ fashion within the OMM (**Figure 6F**, *steps 1 and 2*). Absence of the disulfide-forming cysteines or the entire CHB results in mistargeting of CCDC127 so that it faces the OMM towards the cytosol. The mechanism of OMM insertion remained unclear.

Using KO cells for MTCH2, an OMM insertase (Guna *et al*., 2022), we did not find differences in CCDC127 levels (*data not shown*). However, unassisted membrane insertion of a TMD is conceivable and has previously been demonstrated for other OMM protein, like yeast Fis1 (Kemper *et al*, 2008) and notably also Om45 (Merklinger *et al*, 2012) that is inserted into the OMM from the IMS side, like CCDC127.

CCDC127 is not the only protein with such a topology. Two examples are the yeast proteins Om45 and Mcp3 (Sinzel *et al*., 2016; Song *et al*., 2014; Wenz *et al*., 2014). Both rely on the TIM23 complex for achieving their correct orientation. Mcp3 is guided to the TIM23 complex in the inner membrane of mitochondria (IMM), and is processed by the inner membrane protease (IMP) before becoming inserted by the MIM complex into the OMM. Om45 interacts with the Tim50 receptor of the TIM23 complex in the IMS from where it is shuttled into the OMM by the MIM machinery.

Also, the yeast protein Mix17 contains like CCDC127 disulfide bonds, but seems to rely on the TOM complex for its insertion into the OMM (Resch *et al*., 2025). We thus hypothesize that in general an OMM-single pass transmembrane protein topology with a large domain in the IMS requires the usage of IMM/IMS proteins as “receptors” to achieve correct orientation.

### CCDC127 acts as MICOS partner protein with a function in membrane homeostasis

Our interactome analysis has revealed that the OMM-anchored CCDC127 protein is embedded into a large protein network together with MICOS and SAM complexes that connects OMM and IMM (**Figure 6F**, *step 4*). Such intra-mitochondrial membrane contact sites are particularly important for the structural and functional organization of the IMS. This narrow compartment is composed of a boundary region and the intra-cristae space, both of which are connected via the MICOS complex (Daumke & van der Laan, 2025; Mukherjee *et al*., 2021). Our studies show that CCDC127 is connected to the membrane-bridging MIC60/MIC19 module of MICOS. Moreover, the stable accumulation of CCDC127 and in particular its assembly into higher-molecular weight complexes depend on MICOS. These findings place CCDC127 at the interface between the two functionally and structurally distinct sectors of the IMS. Given the presence of extended coiled-coil domains in CCDC127 is tempting to speculate that the protein may form large oligomeric assemblies that support the spatial organization of the IMS in cooperation with MICOS and other scaffold proteins, like ATAD3A (Arguello *et al*, 2021). The altered cristae morphology and differences in cardiolipin levels of CCDC127-deficient mitochondria together with the structure-function analysis in an accompanying manuscript by Bock-Bierbaum *et al*. as well as findings from a recent correlative light and electron microscopy screen that reported on abnormal mitochondrial morphology and lowered cardiolipin levels upon loss of CCDC127 (Hassdenteufel *et al*., 2025), collectively suggests a role of CCDC127 in membrane architecture and bilayer homeostasis.

## MATERIALS AND METHODS

### Plasmids, cell lines and chemical treatments of cells

For plasmids and cell lines used in this study, see **Tables S1** and **S2**. Cells were cultured in DMEM supplemented with 10% fetal calf serum (FCS) at 37°C under 5% CO_2_.

For the cycloheximide (CHX) chase IP experiments, cells were treated for indicated times with 100 µg/ml CHX (dissolved in DMSO). After each time point the cells were washed with ice-cold PBS-NEM prior to harvesting. For the generation of stable, inducible cell lines the HEK293 cell line–based Flp-In T-REx-293 cell line was used with the Flp-In T-REx system (Invitrogen).

### CRISPR-Cas9-based generation of CCDC127, MIC60 and MIC10 HEK293 knockout cell lines

For the generation of knockout cells, guide RNA Sequence targeting the respective gene was cloned into the pSpCas9(BB)-2A-GFP (PX458) vector, which was a gift from Feng Zhang (Addgene plasmid # 48138) {Ran, 2013 #36The guide RNA sequence (CCDC127: Guide 1:5’-CACCGACGCCGATTTTCTGAGATCATGG-3’ and Guide 2: 5’-CACCGTGAGATCATGGCGTGGTACT-3’, MIC60: Guide 5’-CACCGCAGCATCTCGGTCAAGCGGA-3’ and MIC10: Guide 5’-CACCGAGTCGGAGCTCGGCAGGAAGTGG-3’) was used. HEK Flp-In T-REx-293 or HEK293T cells were transfected with the plasmid using polyethylenimine (PEI; Thermo Fisher Scientific). After 24 h, GFP-positive cells were collected via FACS and single cell clones were seeded onto 96-well plates (1 cell / well). Clonal cell lines were screened for CCDC127, MIC60 or MIC10 protein expression by western blotting.

### Complementation of CCDC127 CRISPR clones

For the complementation of CRISPR clones with different CCDC127 variants, the inducible Flp-In T-REx System was used. All CCDC127 constructs were cloned into the pcDNA5 FRT-TO vector and co-transfected with the pOG44 Vector into the different CRISPR clones by using the transfection reagent FuGene, according to the manufacturer’s guideline. Positive clones were selected with glucose-containing medium (DMEM supplemented with 1 mM sodium pyruvate, 1 x nonessential amino acids, 10% FCS and 500 mg/ml Pen/Strep, 50 µg/ml Uridine) containing 10 µg/ml Blasticidin and 100 µg/ml Hygromycin.

### Generation of MIC10-FLAG expressing HEK293T cells

MIC10-FLAG expressing cells were generated via retroviral transduction of HEK293T ΔMIC10 cells using the pBABE-puro system. The MIC10-FLAG sequence was cloned into the pBABE-puro vector and co-transfected with pGagPol and pVSVG vectors in high-virus-titer-producing HEK293T helper cells using Lipofectamin LTX, according to the manufacturer’s instructions. Virus-containing supernatant from these cells was subsequently used to infect HEK293TΔMIC10 cells. MIC10-FLAG-expressing cells were selected with puromycin and expression was verified by Western blot analysis.

### Acute siRNA-mediated CCDC127 depletion

SiRNA mediated knockdown of CCDC127 was performed by reverse transfection. Therefore 30 pmol RNAi duplex (CCDC127 or Ctr.) were diluted in 500 µl opti-MEM I medium without serum in the well of the plate (6-well). 5 µl Lipofectamine RNAiMAX was added to each well, gently mixed and incubated for 10 min at RT. HEK FLP/IN T-Rex cells were dilute in complete medium without antibiotics that 2.5 ml contains 250,000 cells. Cells were growth for 72h to achieve knockdown of CCDC127.

### Cell proliferation assay to test for growth on different carbon sources

For cell proliferation assay recorded with the cytosmart omni, 15,000 cells were seeded in 48-well dish and incubated at 37°C. For siRNA experiments, cells were seeded after transfection with siRNA as described above. After 48h, the medium was exchanged with DMEM containing galactose (DMEM supplemented with 4.5 g/l galactose, 1 mM sodium pyruvate, 1 x non-essential amino acids, 10% FCS and 50 µg/ml Uridine). Every 6h the coverage of each well was scanned for 6 days using the cytosmart omni.

### Image acquisition

#### Microscope image acquisition

For the image acquisition the microscope LSM 980 with Airyscan 2 and multiplex from Carl Zeiss Microscopy was used with a Plan-Apochromat 63x/1,4 Oil DIC objective and the GaAsP-PMT, Multi-Alkali-PMT detector. The cells were imaged at room temperature with oil as imaging medium. The following fluorochromes were used: Mitotracker CMXRos and Alexa Fluor488. Images were displayed using the acquisition software ZEN 3.3. and were processed using the software OMERO.insight.

#### Western Blot image acquisition

The immunoblotting images were detected using the ChemiDoc Touch Imaging system (Bio-Rad) and an ImageQuant 800 system (Cytiva).

### Analysis of protein complexes by BN-PAGE

Mitochondria analyzed via blue native-PAGE were solubilized on ice in solubilization buffer (1% [w/v] digitonin, 20 mM Tris-HCl, pH 7.4, 0.1 mM EDTA, 50 mM NaCl, 10% [v/v] glycerol, and 1 mM PMSF). After a clarifying spin, loading dye (5 % Coomassie blue G, 500 mM E-amino n-caproic acid in 100 mM Bis Tris pH 7.0) was added and samples applied onto a 4-13% native acrylamide gradient gel and analyzed by Western blot.

### Native and denaturing immunoprecipitation (IP)

To detect protein-protein interactions, native or denaturing IP were performed. Corresponding cell lines were seeded on a 10 cm dish and were grown until confluency of 90%. Before IP was performed, inducible cell lines were induced with doxycycline overnight or for three days. Then the cells were washed with 5 ml ice-cold PBS containing 20 mM NEM and were incubated with another 5 ml ice-cold PBS containing 20 mM NEM for 15 min on ice. The cells were scratched off and pelleted at 700 x g for 3 min at 4°C. For native IP, the pellet was resuspended in 1000 µl native IP buffer (100 mM sodium phosphate pH 8.1, 100 mM sodium chloride, 1% Triton X-100) and was incubated for at least 30 min on ice. For denaturing IP, the pellet was resuspended in 200 µl denaturing IP buffer (30 mM Tris pH 8.1, 100 mM sodium chloride, 5 mM EDTA) and 50 µl 10% SDS was added. The denaturing IP samples were sonicated at maximum amplitude. After cooling down, 750 µl Denaturing buffer with 2.5 % Triton X 100 was added and incubated for at least 30 min on ice. After that, the samples were centrifuged at 21,817 x g for 1 h at 4°C. The supernatant was collected and incubated with 10 µl equilibrated HA beads (monoclonal anti-HA-Agarose produced in mouse) or with Streptactin beads overnight at 4 °C. Then, the beads were washed 4 times with 1 ml washing buffer containing triton X-100 (100 mM sodium phosphate pH 8.1, 1% Triton X-100, 100 mM sodium chloride for native IP; 30 mM Tris pH 8.1, 100 mM sodium chloride, 5 mM EDTA, 1% Triton X-100 for denaturing IP) and 1 x with washing buffer without Triton X-100 with a centrifugation step at 2,000 x g for 1 min at 4°C. After the last centrifugation step, the beads were dried completely and 20 µl reducing Laemmli buffer was added and the samples were boiled for 10 min. The samples were analysed by SDS-PAGE and Western Blot.

For MIC10-FLAG IP 500 µg isolated mitochondria were solubilised for 30 min on ice in 500 µl FIP solubilisation buffer (1% digitonin, 20 mM Tris pH 7.4, 50 mM sodium chloride, 10% glycerol, and 0.1 mM EDTA). After a clarifying spin at 15,000 x g for 10 min at 4°C, supernatant containing solubilised mitochondria were mixed with 20 µl equilibrated FLAG-beads and incubated in a head-over-head shaker at 4°C for 2h. Beads and bound proteins were washed 10 x with FIP-washing buffer (0.3% digitonin, 20 mM Tris pH 7.4, 60 mM sodium chloride, 10% glycerol, and 0.5 mM EDTA) before elution in 75 µl FIP washing buffer containing 100 µg/ml FLAG peptide. Proteins were analysed by SDS-PAGE and Western blot.

### Isolation of crude mitochondria from HEK293 cells

Isolation of crude mitochondria from HEK293 cells was performed as described in (Murschall *et al*, 2021). In short, cells were cultivated for 4 days on 15 cm dishes. For harvesting, the cells were washed 2 times with 10 ml ice-cold 1 x PBS and scraped off using a cell scraper. Afterwards the cells were centrifuged at 500 x g for 5 min at 4°C. The pellets were resuspended in 5 ml 1x M buffer (220 mM mannitol, 70 mM sucrose, 5 mM HEPES-KOH, pH 7.4, 1 mM EGTA-KOH, pH 7.4) containing 1 x Complete TM Protease Inhibitor Cocktail. The cells were homogenized using a precooled potter homogenizer (1,000 rpm, 15 strokes). The homogenized cells were pelleted at 600 x g for 5 min at 4°C. The supernatant containing the crude mitochondria was centrifuged at 8,000 x g for 10 min at 4°C. The pellet was washed with 2 ml ice-cold 1x M buffer (without the Complete^TM^ protease inhibitor cocktail). The crude mitochondria were pelleted at 6,000 x g for 10 min at 4°C and the supernatant was carefully removed. The pellet was resuspended in 400 µl and the concentration was measured using the BCA Reagent ROTI® Quant Assay according to the manufactureŕs instructions.

### Submitochondrial fractionation of isolated mitochondria

For subcellular fraction of CCDC127 40 µg crude mitochondria was centrifuged at 10,000 x g for 5 min at 4°C. Each pellet was resuspended in 95 µl of corresponding buffer (<u>isotonic</u> buffer: 1 x M buffer (220 mM mannitol, 70 mM sucrose 5 mM, HEPES-KOH, pH 7.4, 1 mM EGTA-KOH, pH 7.4); <u>hypotonic</u> buffer: 10 mM HEPES pH 7.4; <u>TX-100</u> buffer: 1x M buffer,1% TX-100; each buffer containing either 40 µg/ml PK or no PK) by pipetting up and down using a 200 µl pipet tip that was cut. The samples were incubated for 5 min at 21 °C and after the incubation time 2.5 µl of 0.2 M PMSF (final concentration 5 mM) was added to all 6 samples and incubation was allowed for a further 5 min on ice. Then, the samples were centrifuged at 10,000 x g for 5 min and the pellets were resuspended with 100 µl of each buffer (isotonic or hypotonic) containing 1 mM PMSF. 4x Laemmli buffer and DTT was added to a final concentration of 50 mM DTT. All samples were boiled at 95°C for 5 min. The samples were analyzed by SDS-PAGE and Western Blot.

### Determination of cellular protein levels by quantitative label-free proteomics

For quantitative label-free proteomics, the experiments were performed as described in (Habich *et al*., 2019). Respective cells were seeded on a 6 well dish. The next day the cells were washed with PBS and harvested by cell scraper and centrifugation at 500 x g for 5 min. The pellets were resuspended in 20 µl lysis buffer (4% SDS in PBS containing protease inhibitor) and were sonicated. Afterwards the samples were boiled for 5 min at 96°C. After cooling down, 80 µl ice-cold acetone was added and stored at -80°C overnight. The next day, TCA precipitations were thawed, and the samples were centrifuged for 15 min at 16.000 x g. The resulting pellets were washed with 500 µl acetone and then air-dried. The pellets were then resuspended in 50 µl 8 M urea in TEAB buffer supplemented with protease inhibitor cocktail and were sonicated. The samples were centrifuged for 15 min at 20,000 x g. The supernatant was transferred into a new tube and the protein concentration of the samples were determined by Pierce Protein Assay Reagent. The assay was performed according to the manufactureŕs protocol and the concentration was measured at 600 nm. 50 µg of each sample was transferred to a new reaction tube and filled up to 40 µl with the Urea/TEAB buffer. Then, DTT with a final concentration of 5 mM was added and the samples were incubated for 1 h at 37°C. Next, chloroacetamide (CAA) with a final concentration of 40 mM was added to the samples and were incubated for 30 min in the dark. For the digest of the peptides, first Lysyl Endopeptidase with an enzyme to substrate ratio of 1:75 was used and the samples were incubated for 4 h at 25°C. For the trypsin digest the samples were first diluted with TEAB buffer to reach a urea concentration below 2 M and then trypsin with an enzyme to substrate ratio of 1:75 was added. The samples were incubated at 25 °C overnight. In the last step, the samples were purified by STAGE tips which were equilibrated with methanol and buffers containing 0.1 % formic acid and 80 % acetonitrile. The samples were loaded on the STAGE tips and were washed with buffers containing 0.1 % formic acid and 80 % acetonitrile. The STAGE tips were completely dried and until measurement stored at 4 °C.

### Determination of protein interaction partners by proteomics (interactome analysis)

For the interactome data an in-gel digest was performed. The native IP was performed as described in the respective section After the native IP was performed the beads were dried and boiled in 20 µl reducing Laemmli buffer (without bromphenolblue) for 10 min. The samples were reduced by addition of DTT with a final concentration of 5 mM and incubated at 56 °C for 30 min. Free cysteine thiols were alkylated by addition of CAA to a final concentration of 40 mM to the samples which were then incubated for 30 min at room temperature in the dark. The samples were run on SDS-PAGE until the samples migrated for 1 cm into the separation gel. Then the gels were fixed for 1 h in fixing solution (10% acetic acid / 20% methanol in water). The gel bands were cut in smaller pieces and were transferred to individual tubes. 100 µl of 50 mM ABC/50% Acetonitrile was added to the gel pieces and were incubated for 20 min at room temperature. The solution was exchanged with fresh 50 mM ABC/50% Acetonitrile and remaining solution was discarded after 20 min incubation. The gel pieces were covered with 100 µl acetonitrile and incubated for 10 min. The gel pieces were then dried in a speedvac for approximately 5 min. A digestion solution of 10 ng/μl of 90 % trypsin and 10 % LysC in 50 mM ammonium bicarbonate (ABC) was added to the gel pieces until the gel pieces were fully covered. The gel pieces were incubated for 30 min at 4°C with the digest solution. After the incubation time, excessive digest solution was removed and 50 mM ABC buffer was used to cover the gel pieces. The samples were incubated overnight at 37°C while shaking at 750 rpm. The next day, the supernatant of the gel pieces were transferred into new tubes. The gel pieces were covered with 100 µl 30% ACN / 3% TFA and incubated for 20 min at room temperature. The extract was combined with the supernatant of the previous step. The gel pieces were covered with 100 µl 100% acetonitrile and again incubated for 20 min at room temperature. The extract was also combined with the supernatant from the previous step and the organic solvents of the samples were reduced in the speedvac until a remaining volume of 50 µl was reached. The samples were acidified by addition of formic acid to a final concentration of 1% and the STAGE tip purification protocol as it is described in the section “Determination of cellular protein levels by quantitative label-free proteomics” was performed. The STAGE tips were stored at 4°C. The mass spectrometry was performed and analysed by the proteomics core facility Cologne.

### Assay to detect redox states of protein thiols

To assess the redox state of CCDC127, 60,000 cells were seeded on a 24-well dish. The cells were cultivated for two days. Before the cells were harvested, they were treated for 19 h with doxycycline (1 µg/ml) to induce CCDC127 expression. The assay was coupled to a CHX treatment. Therefore, the cells were treated with 100 µg/ml CHX before they were harvested and modified. For harvesting the cells were washed with 500 µl ice-cold PB. The PBS was removed and 1000 µl 8% ice-cold TCA was added to each well. The cells were scratched off and transferred to 1.5 ml tubes and were stored at -80°C until the liquid was completely frozen. Samples were thawed at RT and pelleted for 15 min, 13.000 x g, 4°C. The supernatant was removed and 900 µl of ice-cold 5% TCA were added. After vortexing the samples were centrifuged again for 15 min, 13.000xg, 4°C. The TCA was removed completely. Non-reducing Laemmli buffer (2% SDS, 60 mM Tris, pH 6.8, 10% glycerol, 0.0025% bromophenol blue) containing 15 mM mmPEG12 was added to obtain “steady state” samples. The “maximum reduced” and “unmodified” samples contain in addition to Laemmli buffer 10 mM Tris(2-carboxyethyl) phosphine (TCEP) and the alkylating reagent for the “maximum shift” and no reagent for the “unmodified” sample. The maximum reduced samples were boiled for 15 min at 96°C. All samples were sonicated and were analysed by SDS-PAGE and Western Blot.

### Size exclusion chromatography

Analytical size-exclusion chromatography was performed under native conditions to examine the oligomeric state of the endogenous proteins. Cells were washed with 1x PBS, mechanically detached by scraping and sedimented at 500 xg for 5 min. Cell pellets were resuspended in 1 ml native lysis buffer (100 mM sodium phosphate pH 8.0, 100 mM sodium chloride, 1% (v/v) Triton X-100), supplemented with 200 µM PMSF and incubated for 1 h on ice. The lysate was cleared by centrifugation (20 000 xg, 1 h) and loaded on a HiLoad™ 16/600 Superdex 200 preparation grade gel filtration column installed in a liquid chromatography system (Aekta Purifier) from GE Healthcare. A protein size standard was used as a reference, covering a range from 1.35 kDa to 670 kDa (#1511901, Bio-Rad).

### Preparation of Immunofluorescence (IF) samples

To establish the localization of different CCDC127 cysteine mutants, cells were seeded onto poly-L-lysine coated coverslips in 12-well dishes in complete medium. Protein expression was induced by adding 0.1 µg/ml doxycycline for 72 h prior to preparation of IF samples. For Mitotracker staining media was replaced by 1 ml pure media (w/o FCS, P/S) substituted with Mitotracker. Incubation occurred for 1 h at 37 °C. Afterwards the media was replaced by complete media followed by an incubation for 30 min at 37 °C. Cells were washed with prewarmed PBS and 1 ml fixation buffer (4 % PFA in PBS) was added and incubated for 15 min at RT. Cells were washed 3x with PBS and 1 ml fresh blocking buffer (10 mM HEPES pH 7.4, 3% BSA, 0.3 % Triton X-100) was added for 1 h at RT. Cells were washed 3x with PBS. 40 µl per coverslip primary antibody solution in blocking buffer (HA (rabbit) 1:400) was placed onto a parafilm and coverslips were added upside down to the solution for 1 h at RT. Cells were washed 3x with PBS. 40 µl per coverslip secondary antibody solution in blocking buffer (Alexa488 rabbit 1:400) was added onto a parafilm and coverslips were placed upside down to the solution for 1 h at RT in the dark. Cells were washed 3x with PBS and 20 µl prewarmed mixture of Mowiol and DABCO (50 °C) was added onto glass slide and coverslips were placed on top. The samples were dried at 4 °C overnight and sealed with nail polish.

### Fractionation

Cellular fractionation experiments were performed to distinguish between mitochondrial and cytosolic proteins. For this, cells were grown on 15 cm dishes until they reached 80 % confluency. Cells were washed with 10 ml ice-cold PBS and transferred to falcon tubes for centrifugation at 500 x g for 5 min at 4 °C. The cell pellets were resuspended in 2 ml fractionation buffer (20 mM HEPES pH 7.4, 220 mM mannitol, 70 mM sucrose, 1 mM EDTA) and homogenized using the potter with 15 strokes at 1100 rpm. Intact cells and cell debris were pelleted at 800 x g for 5 min at 4 °C. The supernatant was transferred into a 2 ml reaction tube and a total sample (T) was taken. The homogenized sample was centrifuged at 13,000 x g for 15 min at 4 °C to divide a supernatant (cytosolic) and pellet (mitochondrial). The cytosolic fraction was centrifuged at full speed for 10 min and the supernatant was used for TCA precipitation. The mitochondrial pellet was resuspended in 1 ml fractionation buffer and centrifuged at 13,000 x g for 15 min at 4 °C. This washing step was repeated three times before the pellet was resuspended in 80 µl 100 mM Tris/AC pH 8.0. All tree samples (T, cyto, mito) were used for TCA precipitation. Finally, the TCA pellets were resuspended in buffer A and boiled at 96 °C for 5 min.

### Sodium carbonate extraction

In order to access the solubility of mitochondrial proteins, 200 µg freshly isolated mitochondria were centrifuged at 10,000 x g for 5 min and resuspended in 500 µl of ice-cold 0.1 M sodium carbonate solution pH 11.05. Incubation for 1.5 h at 4 °C was performed to open the membranes. A “total (T)” sample was removed. The samples were filled up to 14 ml total volume with sodium carbonate for ultracentrifugation at 90,000 x g for 30 min, 4 °C. The supernatant (14 ml) was collected to perform a TCA precipitation. Both samples (P and SN) were finally resuspended in 131.25 µl water (+ 43.7 µl 4xLaemmli + 8.75 µl 1 M DTT) and boiled at 96 °C for 5 min.

### Oxygen consumption measurements

Mitochondrial respiration was measured with a Seahorse XFe Analyzer (Agilent Technologies). HEK293 cells were seeded onto a 96-well plate after 72h of siRNA-mediated KD of CCDC127 or using a control siRNA. Before the respiration measurements, cells were washed with Seahorse XF DMEM medium enriched with glucose, pyruvate and glutamine and 180 µl fresh medium was added. Cells were incubated for 1 h at 37 °C. The Seahorse injection ports on the sensor cartridge were filled with oligomycin (1 μM), FCCP (1 μM) and Antimycin A/Rotenone (1 μM). Hoechst dye was added in addition to the Antimycin A/Rotenone drug. The sensor cartridges were placed on top of the Seahorse miniplates containing the cells and placed in the Seahorse XFp instrument. Data were analyzed with Wave software (Agilent Technologies). After the run, the plate was placed into a Cytation in order to scan all wells for fluorescence with Hoechst dye. This cell number was used for normalization.

### Transmission Electron Microscopy

Knockdown of CCDC127 was performed 72h prior to TEM sample preparation. HEK293 cells were transferred onto aclar foils. For fixation, the fixative (2 % GA, 2.5 % Sucrose, 3 mM CaCl_2_ in 100 mM HEPES) was prewarmed to 37 °C. The medium was removed carefully and the prewarmed fixative was added carefully. 30 min at RT of incubation followed by 30 m in by 4 °C was performed. The fixative was removed and 100 mM HEPES was added. Samples were stored at 4 °C. Electron microscopy was performed on a JEOL JEM2100PLUSCameraGATAN OneView.

## DATA AVAILABILITY

The datasets generated during and/or analysed during the current study are available from the corresponding author on reasonable request.

## ACKNOWLEDGEMENTS

The Deutsche Forschungsgemeinschaft (DFG, German Research Foundation) funds research in the laboratory of JR through the grants RI2150/5-2 project number 435235019, RTG2550/2 project number 411422114, SPP2453 project number 541742459, CRC1218 - project number 269925409, CRC1678 – project number 520471345 as well as a large instrument grant – project number 533907460. JR also receives funding from the Center for Molecular Medicine (CMMC). The DFG funds research in the laboratory of MvdL through the grants SFB 894/A20 - project number 157660137, FOR 2848/P02 - project number 401510699, SPP 2453/P14 - project number 541758477 and a research program of the Saarland University Medical Faculty (HOMFOR2020) for AvdM; in the laboratory of OD through the grant FOR2848/P06 - project number 401510699. We thank Anja Wittmann, Anika Seiler, Katja Noll and Sibylle Jungbluth for technical support throughout the project, and the CECAD Proteome, Lipidomics and Imaging Facilities for the provision of instrumentation, training, and technical support.

## AUTHOR CONTRIBUTIONS

JR, CZ, MvdL, KvdM designed the study and planned experiments. CZ and AvdM designed, cloned the constructs and generated cell lines. CZ performed bioinformatical analyses. CZ and CECAD proteomics facility carried out the mass spectrometry-based proteomics and performed the bioinformatical analysis of the mass spectrometry data. CZ, RAR, PS, KvdM carried out the biochemical experiments including immunoprecipitation, gel filtration, BN-PAGE, and analytical ultracentrifugation. CZ carried out the fluorescence microscopy experiments and the image analysis. JR, CZ, PS, KvdM, MvdL, TBB and OD analysed the data. JR, CZ, KvdM, MvdL wrote the manuscript with the help and input of all authors.

## REFERENCES

Aaltonen MJ, Friedman JR, Osman C, Salin B, di Rago JP, Nunnari J, Langer T, Tatsuta T (2016) MICOS and phospholipid transfer by Ups2-Mdm35 organize membrane lipid synthesis in mitochondria. J Cell Biol 213: 525–534

Allen S, Balabanidou V, Sideris DP, Lisowsky T, Tokatlidis K (2005) Erv1 mediates the Mia40-dependent protein import pathway and provides a functional link to the respiratory chain by shuttling electrons to cytochrome c. J Mol Biol 353: 937–944

Arguello T, Peralta S, Antonicka H, Gaidosh G, Diaz F, Tu YT, Garcia S, Shiekhattar R, Barrientos A, Moraes CT (2021) ATAD3A has a scaffolding role regulating mitochondria inner membrane structure and protein assembly. Cell Rep 37: 110139

Becker T, Wagner R (2018) Mitochondrial Outer Membrane Channels: Emerging Diversity in Transport Processes. Bioessays 40: e1800013

Bohnert M, Wenz LS, Zerbes RM, Horvath SE, Stroud DA, von der Malsburg K, Muller JM, Oeljeklaus S, Perschil I, Warscheid B et al (2012) Role of mitochondrial inner membrane organizing system in protein biogenesis of the mitochondrial outer membrane. Mol Biol Cell 23: 3948–3956

Brosey CA, Shen R, Tainer JA (2025) NADH-bound AIF activates the mitochondrial CHCHD4/MIA40 chaperone by a substrate-mimicry mechanism. EMBO J

Callegari S, Muller T, Schulz C, Lenz C, Jans DC, Wissel M, Opazo F, Rizzoli SO, Jakobs S, Urlaub H et al (2019) A MICOS-TIM22 Association Promotes Carrier Import into Human Mitochondria. J Mol Biol 431: 2835–2851

Casler JC, Lackner LL (2025) The power of connections: Recent advances in understanding the regulation of mitochondrial dynamics by membrane contact sites. Curr Opin Cell Biol 95: 102535

Chacinska A, Pfannschmidt S, Wiedemann N, Kozjak V, Sanjuan Szklarz LK, Schulze-Specking A, Truscott KN, Guiard B, Meisinger C, Pfanner N (2004) Essential role of Mia40 in import and assembly of mitochondrial intermembrane space proteins. EMBO J 23: 3735–3746

Chen B, Lyssiotis CA, Shah YM (2025) Mitochondria-organelle crosstalk in establishing compartmentalized metabolic homeostasis. Mol Cell 85: 1487–1508

Daumke O, van der Laan M (2025) Molecular machineries shaping the mitochondrial inner membrane. Nat Rev Mol Cell Biol

Diederichs KA, Ni X, Rollauer SE, Botos I, Tan X, King MS, Kunji ERS, Jiang J, Buchanan SK (2020) Structural insight into mitochondrial beta-barrel outer membrane protein biogenesis. Nat Commun 11: 3290

Ding C, Wu Z, Huang L, Wang Y, Xue J, Chen S, Deng Z, Wang L, Song Z, Chen S (2015) Mitofilin and CHCHD6 physically interact with Sam50 to sustain cristae structure. Sci Rep 5: 16064

Doan KN, Grevel A, Martensson CU, Ellenrieder L, Thornton N, Wenz LS, Opalinski L, Guiard B, Pfanner N, Becker T (2020) The Mitochondrial Import Complex MIM Functions as Main Translocase for alpha-Helical Outer Membrane Proteins. Cell Rep 31: 107567

Endo T, Wiedemann N (2025) Molecular machineries and pathways of mitochondrial protein transport. Nat Rev Mol Cell Biol

Fischer M, Horn S, Belkacemi A, Kojer K, Petrungaro C, Habich M, Ali M, Kuttner V, Bien M, Kauff F et al (2013) Protein import and oxidative folding in the mitochondrial intermembrane space of intact mammalian cells. Mol Biol Cell 24: 2160–2170

Fritz S, Rapaport D, Klanner E, Neupert W, Westermann B (2001) Connection of the mitochondrial outer and inner membranes by Fzo1 is critical for organellar fusion. J Cell Biol 152: 683–692

Gabriel K, Milenkovic D, Chacinska A, Muller J, Guiard B, Pfanner N, Meisinger C (2007) Novel mitochondrial intermembrane space proteins as substrates of the MIA import pathway. J Mol Biol 365: 612–620

Ganesan I, Busto JV, Pfanner N, Wiedemann N (2024) Biogenesis of mitochondrial beta-barrel membrane proteins. FEBS Open Bio 14: 1595–1609

Giacomello M, Pyakurel A, Glytsou C, Scorrano L (2020) The cell biology of mitochondrial membrane dynamics. Nat Rev Mol Cell Biol 21: 204–224

Guna A, Stevens TA, Inglis AJ, Replogle JM, Esantsi TK, Muthukumar G, Shaffer KCL, Wang ML, Pogson AN, Jones JJ et al (2022) MTCH2 is a mitochondrial outer membrane protein insertase. Science 378: 317–322

Gupta A, Becker T (2021) Mechanisms and pathways of mitochondrial outer membrane protein biogenesis. Biochim Biophys Acta Bioenerg 1862: 148323

Habich M, Salscheider SL, Murschall LM, Hoehne MN, Fischer M, Schorn F, Petrungaro C, Ali M, Erdogan AJ, Abou-Eid S et al (2019) Vectorial Import via a Metastable Disulfide-Linked Complex Allows for a Quality Control Step and Import by the Mitochondrial Disulfide Relay. Cell Rep 26: 759–774 e755

Hangen E, Feraud O, Lachkar S, Mou H, Doti N, Fimia GM, Lam NV, Zhu C, Godin I, Muller K et al (2015) Interaction between AIF and CHCHD4 Regulates Respiratory Chain Biogenesis. Mol Cell 58: 1001–1014

Harner M, Korner C, Walther D, Mokranjac D, Kaesmacher J, Welsch U, Griffith J, Mann M, Reggiori F, Neupert W (2011) The mitochondrial contact site complex, a determinant of mitochondrial architecture. EMBO J 30: 4356–4370

Helle SC, Kanfer G, Kolar K, Lang A, Michel AH, Kornmann B (2013) Organization and function of membrane contact sites. Biochim Biophys Acta 1833: 2526–2541

Jenkins BC, Neikirk K, Katti P, Claypool SM, Kirabo A, McReynolds MR, Hinton A, Jr. (2024) Mitochondria in disease: changes in shapes and dynamics. Trends Biochem Sci 49: 346–360

Jores T, Klinger A, Gross LE, Kawano S, Flinner N, Duchardt-Ferner E, Wohnert J, Kalbacher H, Endo T, Schleiff E et al (2016) Characterization of the targeting signal in mitochondrial beta-barrel proteins. Nat Commun 7: 12036

Jores T, Lawatscheck J, Beke V, Franz-Wachtel M, Yunoki K, Fitzgerald JC, Macek B, Endo T, Kalbacher H, Buchner J et al (2018) Cytosolic Hsp70 and Hsp40 chaperones enable the biogenesis of mitochondrial beta-barrel proteins. J Cell Biol 217: 3091–3108

Kaurov I, Vancova M, Schimanski B, Cadena LR, Heller J, Bily T, Potesil D, Eichenberger C, Bruce H, Oeljeklaus S et al (2018) The Diverged Trypanosome MICOS Complex as a Hub for Mitochondrial Cristae Shaping and Protein Import. Curr Biol 28: 3393–3407 e3395

Kemper C, Habib SJ, Engl G, Heckmeyer P, Dimmer KS, Rapaport D (2008) Integration of tail-anchored proteins into the mitochondrial outer membrane does not require any known import components. J Cell Sci 121: 1990–1998

Kloppel C, Suzuki Y, Kojer K, Petrungaro C, Longen S, Fiedler S, Keller S, Riemer J (2011) Mia40-dependent oxidation of cysteines in domain I of Ccs1 controls its distribution between mitochondria and the cytosol. Mol Biol Cell 22: 3749–3757

Krimmer T, Rapaport D, Ryan MT, Meisinger C, Kassenbrock CK, Blachly-Dyson E, Forte M, Douglas MG, Neupert W, Nargang FE et al (2001) Biogenesis of porin of the outer mitochondrial membrane involves an import pathway via receptors and the general import pore of the TOM complex. J Cell Biol 152: 289–300

Lauffer S, Mabert K, Czupalla C, Pursche T, Hoflack B, Rodel G, Krause-Buchholz U (2012) Saccharomyces cerevisiae porin pore forms complexes with mitochondrial outer membrane proteins Om14p and Om45p. J Biol Chem 287: 17447–17458

Lionaki E, Aivaliotis M, Pozidis C, Tokatlidis K (2010) The N-terminal shuttle domain of Erv1 determines the affinity for Mia40 and mediates electron transfer to the catalytic Erv1 core in yeast mitochondria. Antioxid Redox Signal 13: 1327–1339

Longen S, Bien M, Bihlmaier K, Kloeppel C, Kauff F, Hammermeister M, Westermann B, Herrmann JM, Riemer J (2009) Systematic analysis of the twin cx(9)c protein family. J Mol Biol 393: 356–368

Merklinger E, Gofman Y, Kedrov A, Driessen AJ, Ben-Tal N, Shai Y, Rapaport D (2012) Membrane integration of a mitochondrial signal-anchored protein does not require additional proteinaceous factors. Biochem J 442: 381–389

Mertins B, Psakis G, Essen LO (2014) Voltage-dependent anion channels: the wizard of the mitochondrial outer membrane. Biol Chem 395: 1435–1442

Mesecke N, Terziyska N, Kozany C, Baumann F, Neupert W, Hell K, Herrmann JM (2005) A disulfide relay system in the intermembrane space of mitochondria that mediates protein import. Cell 121: 1059–1069

Meyer K, Buettner S, Ghezzi D, Zeviani M, Bano D, Nicotera P (2015) Loss of apoptosis-inducing factor critically affects MIA40 function. Cell Death Dis 6: e1814

Michaud M, Gros V, Tardif M, Brugiere S, Ferro M, Prinz WA, Toulmay A, Mathur J, Wozny M, Falconet D et al (2016) AtMic60 Is Involved in Plant Mitochondria Lipid Trafficking and Is Part of a Large Complex. Curr Biol 26: 627–639

Monteiro-Cardoso VF, Rochin L, Arora A, Houcine A, Jaaskelainen E, Kivela AM, Sauvanet C, Le Bars R, Marien E, Dehairs J et al (2022) ORP5/8 and MIB/MICOS link ER-mitochondria and intra-mitochondrial contacts for non-vesicular transport of phosphatidylserine. Cell Rep 40: 111364

Mukherjee I, Ghosh M, Meinecke M (2021) MICOS and the mitochondrial inner membrane morphology - when things get out of shape. FEBS Lett 595: 1159–1183

Murschall LM, Peker E, MacVicar T, Langer T, Riemer J (2021) Protein Import Assay into Mitochondria Isolated from Human Cells. Bio Protoc 11: e4057

Naoe M, Ohwa Y, Ishikawa D, Ohshima C, Nishikawa S, Yamamoto H, Endo T (2004) Identification of Tim40 that mediates protein sorting to the mitochondrial intermembrane space. J Biol Chem 279: 47815–47821

Otera H, Taira Y, Horie C, Suzuki Y, Suzuki H, Setoguchi K, Kato H, Oka T, Mihara K (2007) A novel insertion pathway of mitochondrial outer membrane proteins with multiple transmembrane segments. J Cell Biol 179: 1355–1363

Papic D, Krumpe K, Dukanovic J, Dimmer KS, Rapaport D (2011) Multispan mitochondrial outer membrane protein Ugo1 follows a unique Mim1-dependent import pathway. J Cell Biol 194: 397–405

Paschen SA, Waizenegger T, Stan T, Preuss M, Cyrklaff M, Hell K, Rapaport D, Neupert W (2003) Evolutionary conservation of biogenesis of beta-barrel membrane proteins. Nature 426: 862–866

Petrungaro C, Zimmermann KM, Kuttner V, Fischer M, Dengjel J, Bogeski I, Riemer J (2015) The Ca(2+)-Dependent Release of the Mia40-Induced MICU1-MICU2 Dimer from MCU Regulates Mitochondrial Ca(2+) Uptake. Cell Metab 22: 721–733

Pfanner N, Warscheid B, Wiedemann N (2019) Mitochondrial proteins: from biogenesis to functional networks. Nat Rev Mol Cell Biol 20: 267–284

Rapaport D (2003) Finding the right organelle. Targeting signals in mitochondrial outer-membrane proteins. EMBO Rep 4: 948–952

Resch M, Frickel JS, Dischinger K, Choo RSW, Hell K, Harner ME (2025) The Mia40 substrate Mix17 exposes its N-terminus to the cytosolic side of the mitochondrial outer membrane. J Cell Sci 138

Rothemann RA, Pavlenko E, Mondal M, Gerlich S, Grobushkin P, Mostert S, Racho J, Weiss K, Stobbe D, Stillger K et al (2025) Interaction with AK2A links AIFM1 to cellular energy metabolism. Mol Cell 85: 2550–2566 e2556

Rothemann RA, Stobbe D, Hoehne-Wiechmann MN, Murschall LM, Peker E, Knaup LK, Racho J, Habich M, Gerlich S, Lapacz KJ et al (2024) Interaction with the cysteine-free protein HAX1 expands the substrate specificity and function of MIA40 beyond protein oxidation. FEBS J 291: 5506–5522

Salscheider SL, Gerlich S, Cabrera-Orefice A, Peker E, Rothemann RA, Murschall LM, Finger Y, Szczepanowska K, Ahmadi ZA, Guerrero-Castillo S et al (2022) AIFM1 is a component of the mitochondrial disulfide relay that drives complex I assembly through efficient import of NDUFS5. EMBO J 41: e110784

Schildhauer F, Ryl PSJ, Lauer SM, Lenz S, Barlas AB, Ouzounidis VR, Jeffrey K, Marcu DC, O’Reilly FJ, Graziadei A et al (2025) An NADH-controlled gatekeeper of ATP synthase. Mol Cell 85: 2567–2580 e2512

Shore GC, McBride HM, Millar DG, Steenaart NA, Nguyen M (1995) Import and insertion of proteins into the mitochondrial outer membrane. Eur J Biochem 227: 9–18

Sinzel M, Tan T, Wendling P, Kalbacher H, Ozbalci C, Chelius X, Westermann B, Brugger B, Rapaport D, Dimmer KS (2016) Mcp3 is a novel mitochondrial outer membrane protein that follows a unique IMP-dependent biogenesis pathway. EMBO Rep 17: 965–981

Song J, Tamura Y, Yoshihisa T, Endo T (2014) A novel import route for an N-anchor mitochondrial outer membrane protein aided by the TIM23 complex. EMBO Rep 15: 670–677

Stephan T, Bruser C, Deckers M, Steyer AM, Balzarotti F, Barbot M, Behr TS, Heim G, Hubner W, Ilgen P et al (2020) MICOS assembly controls mitochondrial inner membrane remodeling and crista junction redistribution to mediate cristae formation. EMBO J 39: e104105

Tang J, Zhang K, Dong J, Yan C, Hu C, Ji H, Chen L, Chen S, Zhao H, Song Z (2020) Sam50-Mic19-Mic60 axis determines mitochondrial cristae architecture by mediating mitochondrial outer and inner membrane contact. Cell Death Differ 27: 146–160

Terziyska N, Grumbt B, Bien M, Neupert W, Herrmann JM, Hell K (2007) The sulfhydryl oxidase Erv1 is a substrate of the Mia40-dependent protein translocation pathway. FEBS Lett 581: 1098–1102

Terziyska N, Lutz T, Kozany C, Mokranjac D, Mesecke N, Neupert W, Herrmann JM, Hell K (2005) Mia40, a novel factor for protein import into the intermembrane space of mitochondria is able to bind metal ions. FEBS Lett 579: 179–184

Tiku V, Tan MW, Dikic I (2020) Mitochondrial Functions in Infection and Immunity. Trends Cell Biol 30: 263–275

Varabyova A, Topf U, Kwiatkowska P, Wrobel L, Kaus-Drobek M, Chacinska A (2013) Mia40 and MINOS act in parallel with Ccs1 in the biogenesis of mitochondrial Sod1. FEBS J 280: 4943–4959

Vitali DG, Drwesh L, Cichocki BA, Kolb A, Rapaport D (2020) The Biogenesis of Mitochondrial Outer Membrane Proteins Show Variable Dependence on Import Factors. iScience 23: 100779

von der Malsburg K, Muller JM, Bohnert M, Oeljeklaus S, Kwiatkowska P, Becker T, Loniewska-Lwowska A, Wiese S, Rao S, Milenkovic D et al (2011) Dual role of mitofilin in mitochondrial membrane organization and protein biogenesis. Dev Cell 21: 694–707

Wattenberg B, Lithgow T (2001) Targeting of C-terminal (tail)-anchored proteins: understanding how cytoplasmic activities are anchored to intracellular membranes. Traffic 2: 66–71

Weckbecker D, Longen S, Riemer J, Herrmann JM (2012) Atp23 biogenesis reveals a chaperone-like folding activity of Mia40 in the IMS of mitochondria. EMBO J 31: 4348–4358

Weinhaupl K, Lindau C, Hessel A, Wang Y, Schutze C, Jores T, Melchionda L, Schonfisch B, Kalbacher H, Bersch B et al (2018) Structural Basis of Membrane Protein Chaperoning through the Mitochondrial Intermembrane Space. Cell 175: 1365–1379 e1325

Wenz LS, Opalinski L, Schuler MH, Ellenrieder L, Ieva R, Bottinger L, Qiu J, van der Laan M, Wiedemann N, Guiard B et al (2014) The presequence pathway is involved in protein sorting to the mitochondrial outer membrane. EMBO Rep 15: 678–685

Wiedemann N, Kozjak V, Chacinska A, Schonfisch B, Rospert S, Ryan MT, Pfanner N, Meisinger C (2003) Machinery for protein sorting and assembly in the mitochondrial outer membrane. Nature 424: 565–571

Wrobel L, Trojanowska A, Sztolsztener ME, Chacinska A (2013) Mitochondrial protein import: Mia40 facilitates Tim22 translocation into the inner membrane of mitochondria. Mol Biol Cell 24: 543–554

Xia Y, Zhang Y, Sun Y, He L (2023) CCDC127 regulates lipid droplet homeostasis by enhancing mitochondria-ER contacts. Biochem Biophys Res Commun 683: 149116

Xian H, Liou YC (2021) Functions of outer mitochondrial membrane proteins: mediating the crosstalk between mitochondrial dynamics and mitophagy. Cell Death Differ 28: 827–842

Zhu Y, Akkaya KC, Ruta J, Yokoyama N, Wang C, Ruwolt M, Lima DB, Lehmann M, Liu F (2024) Cross-link assisted spatial proteomics to map sub-organelle proteomes and membrane protein topologies. Nat Commun 15: 3290

Zhuang J, Wang PY, Huang X, Chen X, Kang JG, Hwang PM (2013) Mitochondrial disulfide relay mediates translocation of p53 and partitions its subcellular activity. Proc Natl Acad Sci U S A 110: 17356–17361

